# High early lactational synchrony within baboon groups predicts increased female-female competition and infant mortality

**DOI:** 10.1101/2024.09.09.611196

**Authors:** Jack C. Winans, Niki H. Learn, I. Long’ida Siodi, J. Kinyua Warutere, Elizabeth A. Archie, Jenny Tung, Susan C. Alberts, A. Catherine Markham

## Abstract

Female-female competition may be particularly acute when many females in the same group have dependent young simultaneously, with potential negative consequences for offspring survival. Here, we used 45 years of data on wild baboons (*Papio* sp.) in Amboseli, Kenya, to examine the effects of ‘early lactational synchrony’ (ELS, the proportion of females in a group with an infant <90 days old) on female-female agonistic interactions and infant survival. Because early lactation is energetically demanding for mothers and high-risk for infants, we expected ELS to intensify both female-female aggression and maternal association with males, who can buffer infants from conspecific harassment. In support, when ELS was high, new mothers initiated more agonistic contests. Further, high-ranking females increased their time associating with males, which may limit the ability of low-ranking females to access to male protective services. Finally, high ELS strongly predicted infant mortality. This association may result from both aggression among adult females and infanticidal behavior by peripubertal females. Our findings provide evidence that synchronous reproduction can alter competitive regimes and compromise reproductive outcomes even in nonseasonal breeders.

## Introduction

Intraspecific competition is an important selective force acting on social behavior, morphology, and life history^1–3^. In many vertebrates, females compete for food, mates, social partners, and other resources, with tactics ranging from mating interference to infanticide^4–10^. Female-female competition can have important fitness effects, as indicated by the association between high dominance rank or increased aggression and higher female reproductive success in many species^11–16^. For group-living species, group demography is expected to shape the dynamics of female-female competition. Specifically, because female reproduction is energetically costly and food resources are often limited, a group’s size, age structure, sex ratio, and reproductive activity can affect the level of female competition and its fitness consequences^17–28^. Female competition may thus be a mediating mechanism through which group-level social structure affects individual fitness.

Among social mammals, female reproductive state is a particularly salient source of variation in competitive landscapes because a female’s resource and social requirements depend on whether she is cycling, pregnant, lactating, or reproductively quiescent^29^. Accordingly, shared requirements can promote association by females in similar reproductive states^30^, increase competition for reproductive state-specific resources^5^, or both. The importance of reproductive state in shaping female-female competition is supported by the observation that conspicuous female-female competition is more frequent among species that exhibit high reproductive synchrony^3,5^. For example, female topi antelopes (*Damaliscus lunatus*), red deer (*Cervus elaphus*), feral horses (*Equus caballus*), and golden snub-nosed monkeys (*Rhinopithecus roxellana*) display increased rates of aggression and/or disruption of copulations involving female competitors during their short breeding seasons^31–34^. While birth synchrony may confer benefits (e.g. predator swamping^35^; matching reproductive timing to resource availability^36^), distributing births throughout the year may reduce female reproductive competition. However, even in nonseasonal breeders, high reproductive synchrony can sometimes occur. In these species, the behavioral and fitness consequences of competition resulting from reproductive synchrony remain poorly understood. Addressing this gap is important for understanding the relationship between resource availability, competition, and reproductive timing, which can vary even between closely related, ecologically similar species^37,38^. Furthermore, it is well-recognized that interindividual variation (e.g., in reproductive state) affects the social interactions that produce variation in group-level competitive landscapes, affiliative opportunities, and dominance hierarchies)^39,40^. However, little is known about how group-level variation in social structure affects individual fitness in social mammals.

In this study, we investigated how female-female competition during lactation affects female behavior and infant survival in wild baboons (*Papio* sp.). We hypothesized that female-female competition would intensify in baboon social groups with high ‘early lactational synchrony’ (i.e., a high proportion of females with infants <90 days old) compared to groups with low early lactational synchrony, and that intensified competition would lead to negative fitness-related outcomes for mothers and their dependent offspring. We expected that group-level synchrony in early lactation (as opposed to synchrony in other stages of the interbirth interval) would be particularly consequential for female behavior and infant survival. This expectation stems from the unique dual challenge faced by female baboons in early lactation: first, increased energetic demands, coupled with clinging infants that can compromise foraging efficiency^41^; and second, the need to protect their vulnerable infants from harassment and infanticide by conspecifics^42–45^. These challenges are interconnected, as some harassment-prevention strategies (e.g., social vigilance^46^, ‘restrictive mothering’^41^) may constrain food intake. As a result, females with neonates compete for both food^47–49^ and adult male associates, who can help buffer mothers and their infants from conspecific harassment and infanticide^6,50–55^.

We focused on lactational synchrony in the early stage of infancy because, while the energetic demands of lactation and infant transport on mammalian mothers^42,56,57^ typically increase as infants grow^42,43^, maternal energy balance does not linearly track the underlying demands of lactation^58–60^. In particular, as infant movement becomes increasingly independent from mothers, infants are less likely to impede a mother’s ability to ingest food at rates that compensate for the energetic demands of the postpartum period^46,61,62^. In baboons, for example, mothers carry their offspring ventrally during early lactation^41^, introducing two potential forms of feeding interference that are mitigated in later lactation: active disruption of maternal food intake by infants, and passive interference when infants become fatigued from ventral clinging and mothers must sit to support them^41^. Both forms can be costly, as suggested by the tendency of mothers in later lactation to rebuff infants when they approach while mothers are feeding^41^, and by the increased time early lactating mothers spend seated while feeding^41^, which limits their ability to access food outside an arm’s reach.

Beyond their energetic needs, new mothers must also simultaneously defend against aggression towards their neonates, which is common across mammals^7,63–65^. In the baboons we investigated here and in several related baboon species, one form of defense involves protective ‘primary associations’ with specific males (who are sometimes their infants’ fathers)^52,66,67^. The presence of these males in early life has both immediate and lasting impacts on offspring welfare, foraging success, and survival^52,55,68,69^. Because female baboons compete for male social partners^6,53^, high early lactational synchrony should intensify female competition over primary associate males. This form of competition likely explains why lactating chacma baboons (*Papio ursinus*) direct aggression towards fertile females who mate with their primary associate male^6^, and why female-female aggression intensifies with high dyadic and group-level reproductive state synchrony^50^. In the absence of protector males, low-ranking females may increase their reliance on other counterstrategies to infanticide, such as ‘restrictive mothering’ (i.e., limiting opportunities for infants to break physical contact) and increased social vigilance^41^. Given other costs to maternal foraging efficiency during early lactation^41^, increased reliance on these alternative counterstrategies to infanticide in periods of high lactational synchrony may constrain the energy budgets of some mothers even further. Data on chacma baboons also support this idea: low-ranking mothers exhibit longer interbirth intervals after giving birth in close temporal proximity to other groupmates^37^. Thus, we reasoned that high group-level synchrony in early lactation constrains the ability of female baboons to flexibly compensate for the high energetic demands of lactation.

Our study subjects were members of multiple multi-male, multi-female social groups in a wild baboon population (*P. cynocephalus* with significant *P. anubis* admixture^70,71^) in the Amboseli basin of southern Kenya^72^. Females in this population ‘inherit’ their dominance rank matrilineally and give birth year-round, almost always to a single infant^73–75^. We list the predictions of our hypothesis that high early lactational synchrony increases female-female competition and infant mortality risk in Table 1. To test these predictions, we analyzed female-female agonistic interaction rates, time spent on competition-related behaviors by females, and infant mortality risk as functions of early lactational synchrony. We also examined the most prevalent sources of infant mortality during periods of high early lactational synchrony to gain insight into the specific risks imposed by female-female competition. These analyses provide an important example of how group-level processes can affect individual fitness, as well as insights into the expression and outcomes of female-female competition in vertebrates, a topic of increasing interest in animal behavior in the past two decades^4,5,76,77^.

**Table 1.** Hypotheses and predictions. Predictions stemming from our main hypothesis that high ‘early lactational synchrony’ within baboon groups (the proportion of adult females with an infant <90 days old) increases female-female competition and leads to negative fitness outcomes.

| Sub-hypothesis | Prediction and rationale | Supported by this study? |
| --- | --- | --- |
| 1. Baboons in early lactation are focal points for female-female competition | 1.1. Direct competition: Females in early lactation experience agonistic interactions with other adult females more frequently than adult females in other reproductive states, because they have a heightened interest in maintaining proximity to males and a reduced ability to meet their energetic demands with sufficient food intake rates. | Yes |
|  | 1.2. Competition-related behavior: Females in early lactation spend more of their feeding time in ventral contact with an infant than adult females in other reproductive states, leading to foraging constraints during an energetically demanding time. | Yes |
|  | 1.3. Competition-related behavior: Females in early lactation spend less time foraging than females in other reproductive states, because social vigilance constrains their time budgets and ventral infant contact constrains their physical mobility. | Yes |
|  | 1.4. Competition-related behavior: Females in early lactation spend more time in close proximity to adult males than adult females in other reproductive states (with the exception of females with turgid sexual swellings), because primary male associates can buffer against conspecific threats. | Yes |
| 2. Female-female competition intensifies when early lactational synchrony is high | 2.1. Direct competition: Early lactating females initiate agonistic interactions more frequently when early lactational synchrony is higher, because more of their groupmates share a heightened interest in maintaining proximity to males and a reduced ability to meet their energetic demands. | Yes |
|  | 2.2. Competition-related behavior: As early lactational synchrony increases, high-ranking females spend more time in close proximity to adult males and low-ranking females spend less time in close proximity to males, because high-ranking females aggressively exclude low-ranking females from associating with males. | Yes |
|  | 2.3. Competition-related behavior: As early lactational synchrony increases, low-ranking females spend more time foraging, but high-ranking females do not, because agonistic behavior directed towards low-ranking females constrains their ability to forage efficiently. | No/Partial |
|  | 2.4. Competition-related behavior: As early lactational synchrony increases, low-ranking females spend more of their feeding time in ventral contact with an infant, but high-ranking females do not, because low-ranking females are excluded from associating with males and increase reliance on 'restrictive' mothering to protect their infants. | Yes |
| 3. Female-female competition related to high early lactational synchrony increases infant mortality risk | 3.1. Fitness consequence of competition: Infant mortality risk is higher on days with higher early lactational synchrony, because maternal and infant condition are compromised by the social and energetic costs of increased female-female competition. | Yes |
|  | 3.2. Competition-related behavior/direct competition: Infanticide is a more common cause of infant death on days with higher early lactational synchrony, because more mothers are excluded from a typical counter-infanticide strategy (formation of primary associations with males). | Yes |
|  | 3.3. Direct competition: Infanticides on days with high early lactational synchrony are performed by adult females, because high early lactational synchrony increases reproductive competition between adult females. | No |

## Results

### Characterizing early lactational synchrony across the study period

We began by measuring the proportion of females in each social group who were in early lactation on each day across all group-days of observation from November 1976 to December 2021. We defined early lactation as the first 90 days of postpartum amenorrhea when a mother had a living infant. In this period, infants are dependent on their mothers for almost all nutrition and transport^41–43,78^. We based this 90-day threshold on careful consideration of previous research on changes in maternal behaviors related to the ‘dual challenge’ of protecting vulnerable infants while meeting rising energetic demands^41,42^. In support of our focus on the first 90 days postpartum, females in this period engaged in higher rates of agonism and spent more time carrying their infant ventrally, less time foraging, and more time close to adult males, compared to later stages of postpartum amenorrhea (Figs. S1-S2). We also performed parallel analyses for later stages of postpartum amenorrhea to test if group-level synchrony in any of these stages was also correlated with behavioral indicators of female-female competition or with infant mortality. We found no evidence that synchrony in a later stage of postpartum amenorrhea was more salient than the 90-day postpartum window we focus on here (Table S1 summarizes all statistical models and model results are given in the Supplementary Data file; models 34-39 analyzed these alternative metrics of lactational synchrony).

The proportion of females in early lactation varied considerably across the study period, from group-days on which no females (0.000) were in early lactation to group-days on which more than 70% (0.714) of females were in early lactation (Fig. S3). The mean proportion of females in early lactation on a given group-day was 0.121 ± 0.102 s.d. (median: 0.105; Fig. S3a) and, on average, groups contained 1.958 ± 1.739 s.d. total females in early lactation on any given day (median: two females, Fig. S3b). The number of adult females per group ranged from three to 33.

While daily early lactational synchrony was low on average, periods of high early lactational synchrony were not uncommon. Across the 138 group-years in our 1976-2021 dataset where group observations were uninterrupted (e.g., the groups were not undergoing a permanent fission or fusion and/or were consistently observed across the full year), the mean maximum value of early lactational synchrony per year was 0.320 ± 0.087 s.d. This estimate, which corresponds to the 95^th^ percentile value over all group-days, suggests that periods of high early lactational synchrony are a recurring feature of the system (even though they are typically brief in duration: Table S2). Therefore, we used 0.320 as a benchmark of high early lactational synchrony that was met with some regularity (hereafter, ‘high synchrony benchmark’). Under this definition, groups experienced periods of high early lactational synchrony at a mean frequency of 0.665 times per year ± 0.441 s.d. (or once every 1.504 years). On average, periods of high early lactational synchrony lasted 28.058 days ± 19.120 s.d.

### Baboons in early lactation were focal points for female-female competition

To test whether early lactation corresponded to a period of intense competition for individual female baboons (Table 1, sub-hypothesis 1), we tested for changes in behaviors plausibly related to competition over food or males across different stages of the female interbirth interval. Using generalized linear mixed models (GLMMs) fit to behavioral data on 270 adult females (60,665 focal animal samples collected in 14 social groups from 1999-2021), we found that females in early lactation were involved in agonistic interactions (as either initiators or recipients) with other adult females at a higher rate than were females in any other reproductive state (Fig. 1a; prediction 1.1; model 1). For example, the average early lactating female was predicted to experience an agonistic interaction with another adult female groupmate at a rate of 0.350 acts/hour, compared to 0.143 to 0.270 acts/hour for females in other reproductive states. Females in early lactation also spent more of their feeding time in ventral contact with an infant (Fig. 1b; prediction 1.2; model 2) and less time foraging (Fig. 1c; prediction 1.3; model 3) than did females at any other reproductive state we considered. Finally, females in early lactation spent more time within 5 m of at least one adult male than females in any other reproductive state (Fig. 1d; prediction 1.4; model 4), with the exception of females with turgescent sexual swellings, who are highly attractive to adult males.

**Figure 1.**
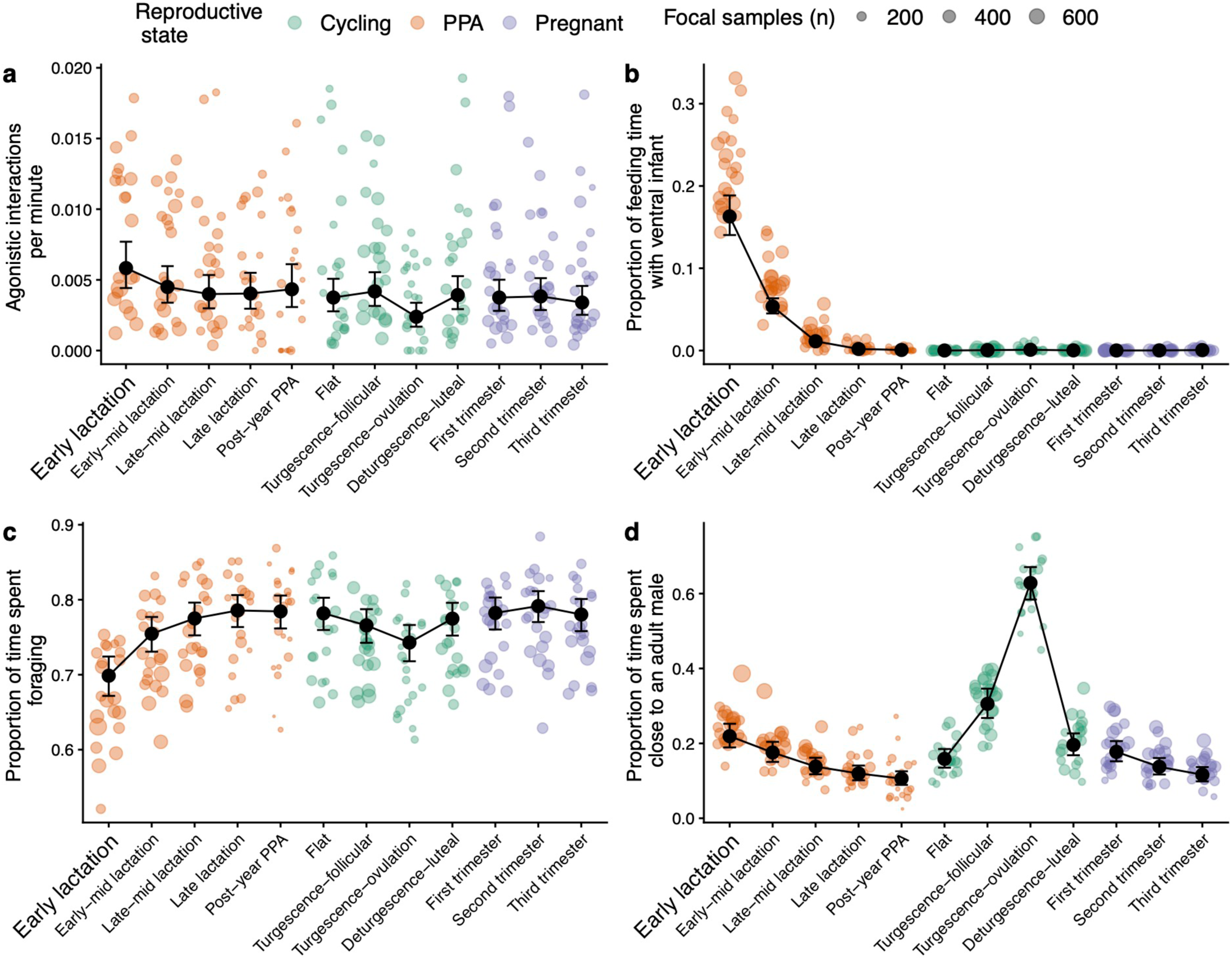
Changes in competition-related behavior across the interbirth interval. Among adult females, (a) the mean rate of agonistic interactions with other adult females, (b) the mean proportion of feeding time spent in ventral contact with an infant, (c) the mean proportion of time spent foraging, and (d) the mean proportion of time spent within 5 m of an adult male varied both between and within the three ‘major’ reproductive states (postpartum amenorrhea (PPA), in orange, ovarian cycling in green, and pregnancy in purple). ‘Minor’ reproductive states are indicated on the x-axis, where early lactation is indicated with enlarged text. Within PPA, early lactation shows a trough in time spent foraging (panel c) and peaks in rate of agonistic interactions (panel a), feeding time spent ventrally supporting an infant (panel b), and time spent in proximity to males (panel d). Black points represent the GLMM-predicted values of each response variable for each reproductive state (holding other fixed effects constant at their observed mean value) and error bars represent 95% confidence intervals of these model predictions. Each colored circle represents the mean observed value for females in a given minor reproductive state in a complete hydrological year. These circles are proportionally sized to the number of focal samples that were conducted on females in that reproductive state and year. For (a), (c), and (d), n = 60,665 focal samples and for (b), n = 45,706 focal samples. In all four panels, the differences between early lactating females and females in all other reproductive states were statistically significant (see Supplementary Data file for model results). See Supplementary Methods for definitions of the reproductive states.

The higher rate of female-female agonistic interactions experienced by early lactating females was most strongly driven by two sets of interactions: (i) agonistic behavior initiated by early lactating females toward other early lactating females (Fig. 2a; model 5), and (ii) agonistic behavior received by early lactating females from pregnant females (Fig. 2b; model 6). Controlling for the number and reproductive state of potential interaction partners, early lactating females initiated agonistic behavior toward other early lactating females at a rate that was 1.8x higher than the rate initiated by post-early lactation postpartum amenorrhea females (β = −0.603, 95% CI: [−1.072, −0.135]) and 1.8x higher than the rate initiated by cycling females (β = −0.573, 95% CI: [−1.015, −0.132]). The rate at which early lactating females initiated agonistic interactions towards other early lactating females was not significantly different from the initiation rate for pregnant females (β = −0.002, 95% CI: [−0.419, 0.415]).

**Figure 2.**
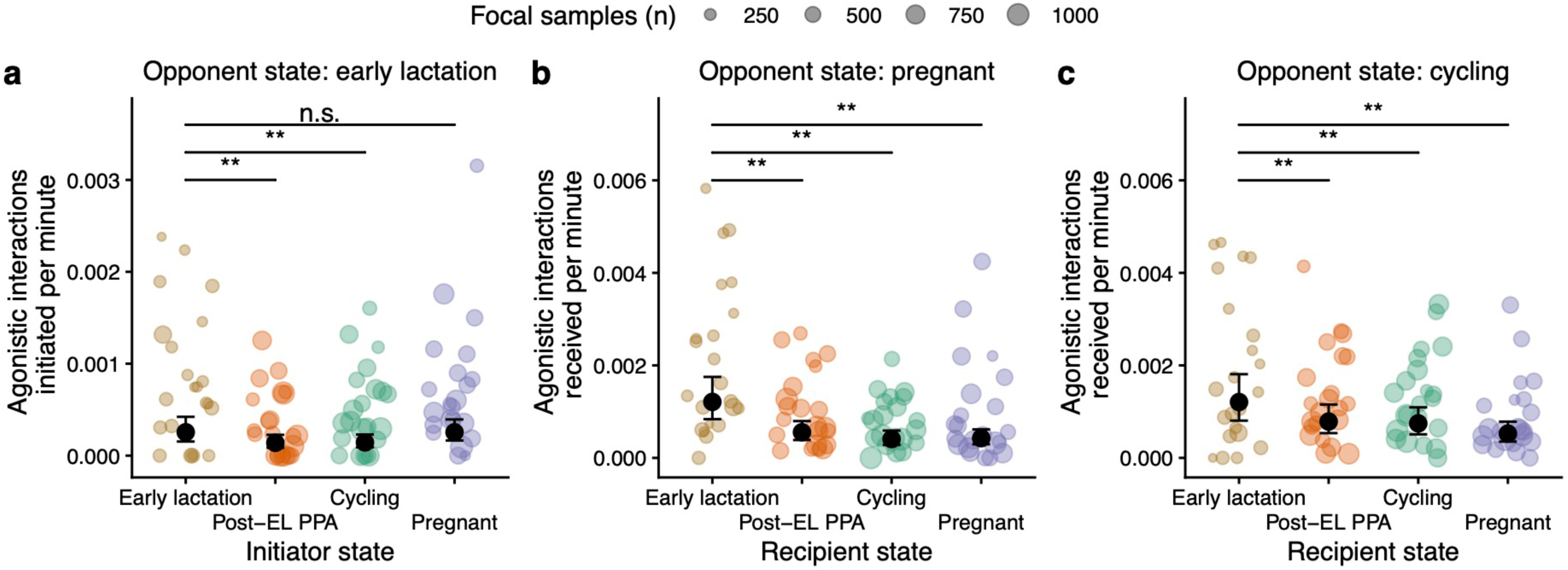
Dyadic agonistic interaction rates as a function of the reproductive states of the interacting females. Each panel depicts the rate per minute of agonistic interactions either (a) initiated toward, or (b-c) received from females of a particular reproductive state as function of the reproductive state of the initiator (a) or recipient (b-c). (a) depicts the rate of agonistic interactions initiated toward females in early lactation, while (b) and (c) depict the rates of agonistic interactions received from pregnant and cycling females, respectively. ‘Post-EL PPA’ refers to females who were in postpartum amenorrhea, but were no longer in early lactation. In each panel, the black circles represent the GLMM-predicted values of each response variable for each reproductive state (holding other fixed effects constant at their observed mean value) and the error bars represent 95% confidence intervals of these model predictions. The colored circles represent the mean observed value for females in a given reproductive state in a complete hydrological year, and these circles are sized proportionally to the number of focal samples that were conducted on females of that reproductive state in that year. For (a), n = 48,335 focal samples, for (b), n = 58,602 focal samples, and for (c), n = 59,679 focal samples. In each model, the reference category was early lactation. ‘**’ indicates a statistically significant difference in the rate of agonistic interactions between early lactating females and females in one of the other reproductive states, while ‘n.s.’ indicates that the difference in agonistic interaction rate between females in early lactation and females in one of the other reproductive states was not statistically significant. We considered any fixed effect to be statistically significant when the 95% confidence interval of the coefficient estimate did not overlap zero. Each of these models controlled for variation in the number of potential opponents in the reproductive state of interest. Note that the scale of the y-axis in (b) and (c) is double that of the y-axis in (a), because, on average, females received agonistic interactions at a higher rate than they initiated agonistic interactions (see Fig. S8).

Early lactating females received agonistic interactions from pregnant females at a rate that was 2.2x higher than the rate received by post-early lactation postpartum amenorrhea females (β = −0.781, 95% CI: [−1.058, −0.504]), 3.0x higher than the rate received by cycling females (β = −1.090, 95% CI: [−1.377, −0.803]), and 2.8x higher than the rate received by other pregnant females (β = −1.044, 95% CI: [−1.327, −0.760]). Cycling females seemed to target early lactating females less specifically than pregnant females did, but still posed an outsized threat (Fig. 2c; model 7): early lactating females received agonistic interactions from cycling females at a rate that was 1.5x higher than the rate received by postpartum females after the end of early lactation (β = −0.429, 95% CI: [−0.691, −0.168]), 1.6x higher than the rate received by cycling females (β = −0.482, 95% CI: [−0.740, −0.223]), and 2.3x higher than the rate received by pregnant females (β = −0.832, 95% CI: [−1.113, −0.551]). No other combinations of female reproductive states exhibited significant differences in the rates of agonistic interactions initiated or received (models 8-12).

### Female-female competition intensified when early lactational synchrony was high

To determine how group-level synchrony in early lactation was related to the intensity of within-group female-female competition (Table 1, sub-hypothesis 2), we used focal sample data collected from 1999-2021 to operationalize several behavioral proxies of competition (n = 235 early lactating females and 7,640 focal samples). First, we measured how rates of agonistic interactions initiated by early lactating females varied as a function of early lactational synchrony (i.e., metrics of direct contests). Then, we tested for rank-related differences in the relationship between early lactational synchrony and several behavioral metrics of female competition: time spent with adult males, time spent foraging, and feeding time spent in ventral contact with an infant. We also examined some competition-related behaviors in pregnant females (n = 250 pregnant females and 16,991 focal samples), because of the outsized agonistic threat they posed to early lactating females (Fig. 2b).

We found that early lactating females initiated agonistic interactions more frequently as early lactational synchrony increased, controlling for the number of other adult females present in the same group (Table 2; Fig 3a; prediction 2.1; model 13; β = 0.186, 95% CI: [0.009, 0.363]). On days when a group was at our high synchrony benchmark (32.0% of females in early lactation), an early lactating female was predicted to initiate agonistic interactions with other adult females at a rate 1.4x higher than on days when the group was at the population median value of early lactational synchrony (10.5% of females in early lactation).

**Figure 3.**
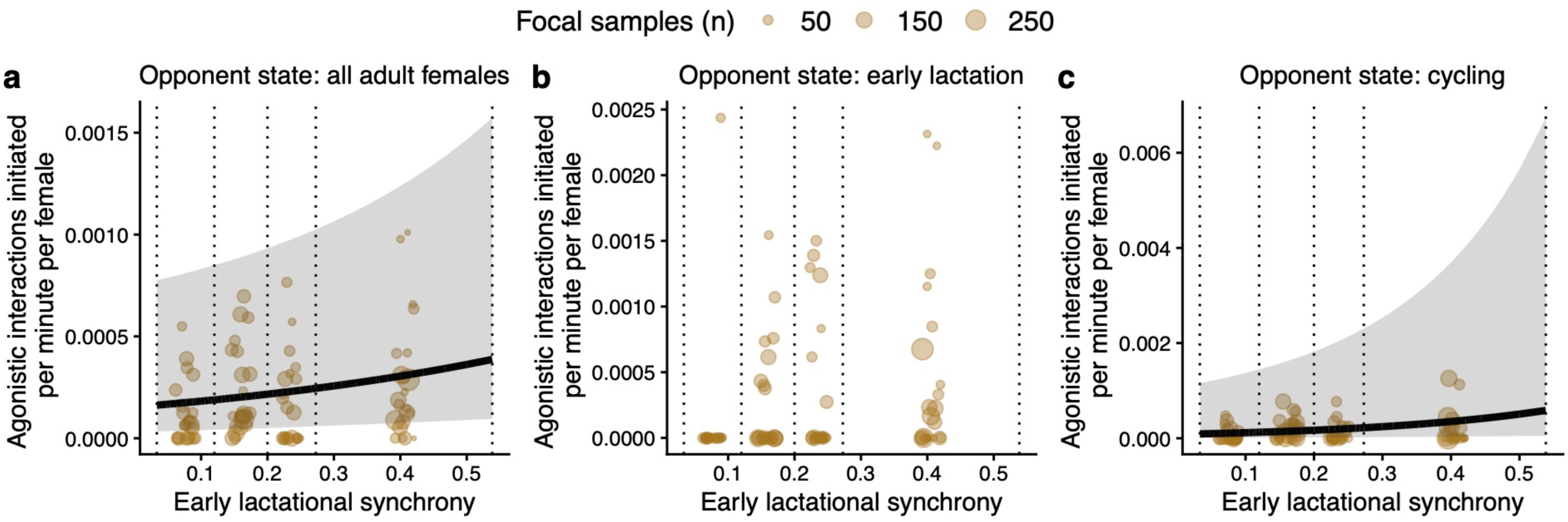
Rates of agonistic interactions initiated by females in early lactation as a function of early lactational synchrony. These panels depict rates per minute of agonistic interactions initiated by females in early lactation toward (a) all adult females, (b) other females in early lactation, and (c) cycling females as functions of early lactational synchrony. These rates are divided by the number of (a) nonfocal adult females, (b) nonfocal early lactating females, or (c) nonfocal cycling females, such that these rates indicate agonistic interactions per minute per potential interaction partner. In all panels, each brown circle represents the mean observed rate calculated from all data collected on days that were within a given quartile of early lactational synchrony in a complete hydrological year. These circles are proportionally scaled in size according to the number of focal samples contributing to the mean rate. The vertical dotted lines indicate the bounds of the quartiles of early lactational synchrony. For (a), n = 7,640 focal samples, for (b) n = 6,248 focal samples, and for (c), n = 7,536 focal samples. The solid black lines in (a) and (c) correspond to the model-predicted relationships between early lactational synchrony and the per-female agonistic interaction rate (holding other fixed effects constant at their observed mean value), and the shaded regions depict the 95% confidence intervals of these predicted relationships. Model fits are only depicted in cases where early lactational synchrony had a significant relationship with the response variable (i.e., the 95% confidence interval of the coefficient estimate for early lactational synchrony did not cross zero). While early lactating females initiated more agonistic interactions overall when synchrony was higher (a), (b) suggests they did not increase rates of agonism with individual early lactating females. Meanwhile, (c) suggests that early lactating females increased rates of agonism with individual cycling females when synchrony was higher.

**Table 2.** Agonistic interaction initiation and time spent with males are predicted by early lactational synchrony. Model-averaged estimates for the best-fitting models (ΔAIC_c_ < 2) predicting the rate at which early lactating females initiated agonistic interactions toward other adult females (n = 7,640 focal samples of 235 females) and the proportion of time that early lactating females spent in close proximity to males (n = 7,640 focal samples of 235 early lactating females). The top 20 candidate models predicting these response variables can be found in the Supplementary Data file. Bold effects are those where the 95% CI does not overlap zero. Model-predicted values are presented to aid interpretation of estimates and were calculated for the extreme values of each significant fixed effect, while holding values for other fixed effects constant at their observed mean value.

| Response variable | Fixed effect | Estimate | Std. Error | 95 % CI | Model-predicted values |
| --- | --- | --- | --- | --- | --- |
| Rate of agonistic interactions initiated toward adult females (model 13) | **(intercept)** | **-6.558** | **0.181** | **[-6.913, -6.203]** | – |
|  | **Early lactational synchrony** | **0.186** | **0.090** | **[0.009, 0.363]** | 0.033 (min) = 0.063 acts/h<br>0.538 (max) = 0.150 acts/h |
|  | **Group size\*** | **-0.690** | **0.282** | **[-1.243, -0.137]** | 13 members (min) = 0.340 acts/h<br>117 members (max) = 0.010 acts/h |
|  | **Dominance rank** | **0.940** | **0.107** | **[0.731, 1.149]** | Rank 0.0 (proportional, lowest) = 0.018 acts/h<br>Rank 1.0 (proportional, highest) = 0.366 acts/h |
|  | Age | 0.037 | 0.070 | [-0.101, 0.174] | – |
|  | **# of adult females present** | **0.781** | **0.292** | **[0.208, 1.354]** | 3 females (min) = 0.017 acts/h<br>30 females (max) = 0.585 acts/h |
| Proportion of time spent close to an adult male (model 18) | **(intercept)** | **-1.397** | **0.115** | **[-1.624, -1.171]** | – |
|  | **Dominance rank** | **-0.013** | **0.005** | **[-0.024, -0.003]** | – |
|  | Early lactational synchrony | 0.301 | 0.163 | [-0.019, 0.621] | – |
|  | **Dominance rank x Early lactational synchrony** | **-0.059** | **0.016** | **[-0.090, -0.027]** | Rank 29 (ordinal, lowest), ELS 0.033 (min) = 0.148<br>Rank 29 (ordinal, lowest), ELS 0.538 (max) = 0.074<br>Rank 1 (ordinal, highest), ELS 0.033 (min) = 0.259<br>Rank 1 (ordinal, highest), ELS 0.538 (max) = 0.283 |
|  | **# of adult males present** | **0.038** | **0.004** | **[0.030, 0.046]** | 1 male (min) = 0.169<br>30 males (max) = 0.380 |
|  | **# of cycling females present** | **-0.017** | **0.005** | **[-0.027, -0.007]** | 0 females (min) = 0.246<br>20 females (max) = 0.190 |
| | Rainfall | $3.5 \times 10^{-4}$ | $3.0 \times 10^{-4}$ | $[-2.4 \times 10^{-4}, 0.001]$ | – |
|  | Age | -0.001 | 0.003 | [-0.007, 0.005] | – |

\*Because this model controlled for the number of nonfocal adult females present, the negative effect of group size should be interpreted such that females in larger groups initiated fewer agonistic interactions *per potential female agonistic partner*, not that females in larger groups initiated fewer agonistic interactions overall (e.g., see Fig. S8).

Two changes to female-female interaction patterns could explain this relationship. First, because the most common targets of aggression from early lactating females were other early lactating females (Fig. 2a), increased aggression on days of high early lactational synchrony could arise from the greater abundance of targets. In support of this possibility, when controlling for the number of other early lactating females present, early lactational synchrony no longer predicted the rate at which early lactating females initiated agonistic interactions with other early lactating females (Fig. 3b; model 14; β = 0.157, 95% CI: [−0.376, 0.690]). This result suggests that females in early lactation simply have more new mother competitors on days when synchrony is high. A second, non-mutually exclusive possibility is that early lactating females also increased the rates at which they targeted specific individuals. This explanation was also partially supported: while rates of agonistic interactions within dyads of early lactating females did not increase with increasing early lactational synchrony (Fig. 3b), early lactating females did increase the rate at which they targeted individual cycling females (controlling for the number of cycling females present: Fig. 3c; model 15; β = 0.397, 95% CI: [0.035, 0.759]). Thus, higher agonistic interaction rates in periods of high early lactational synchrony also partially resulted from increased aggression between early lactating and cycling females. We found no evidence that early lactational synchrony predicted rates of agonistic interactions initiated by early lactating females toward individual pregnant females (model 16; β = −0.177, 95% CI: [−0.671, 0.316]) or individual females in postpartum amenorrhea who were no longer in early lactation (model 17; β = −0.295, 95% CI: [−0.608, 0.219]).

If high early lactational synchrony intensifies female-female competition over male social partners, high-ranking females should be most successful at staying close to adult males because of their ability to displace lower status females (prediction 2.2). In support of this prediction, we found that the proportion of time that both early lactating females and pregnant females spent within 5 m of an adult male was predicted by their dominance rank. Higher-ranking females spent more time in close proximity to males than lower-ranking females, and the magnitude of this rank difference increased with higher group-level early lactational synchrony (Table 2; Fig. 4a,b; model 18, GLMM for early lactating females: ordinal dominance rank: β = −0.013, 95% CI: [−0.024, −0.003]; early lactational synchrony: β = 0.301, 95% CI: [−0.019, 0.621]; interaction term: β = −0.059, 95% CI: [−0.090, −0.027]; model 19, GLMM for pregnant females: ordinal dominance rank: β = −0.079, 95% CI: [−0.015, −0.001]; early lactational synchrony: β = 0.413, 95% CI: [0.185, 0.612]; interaction term: β = −0.053, 95% CI: [−0.080, −0.027]).

**Figure 4.**
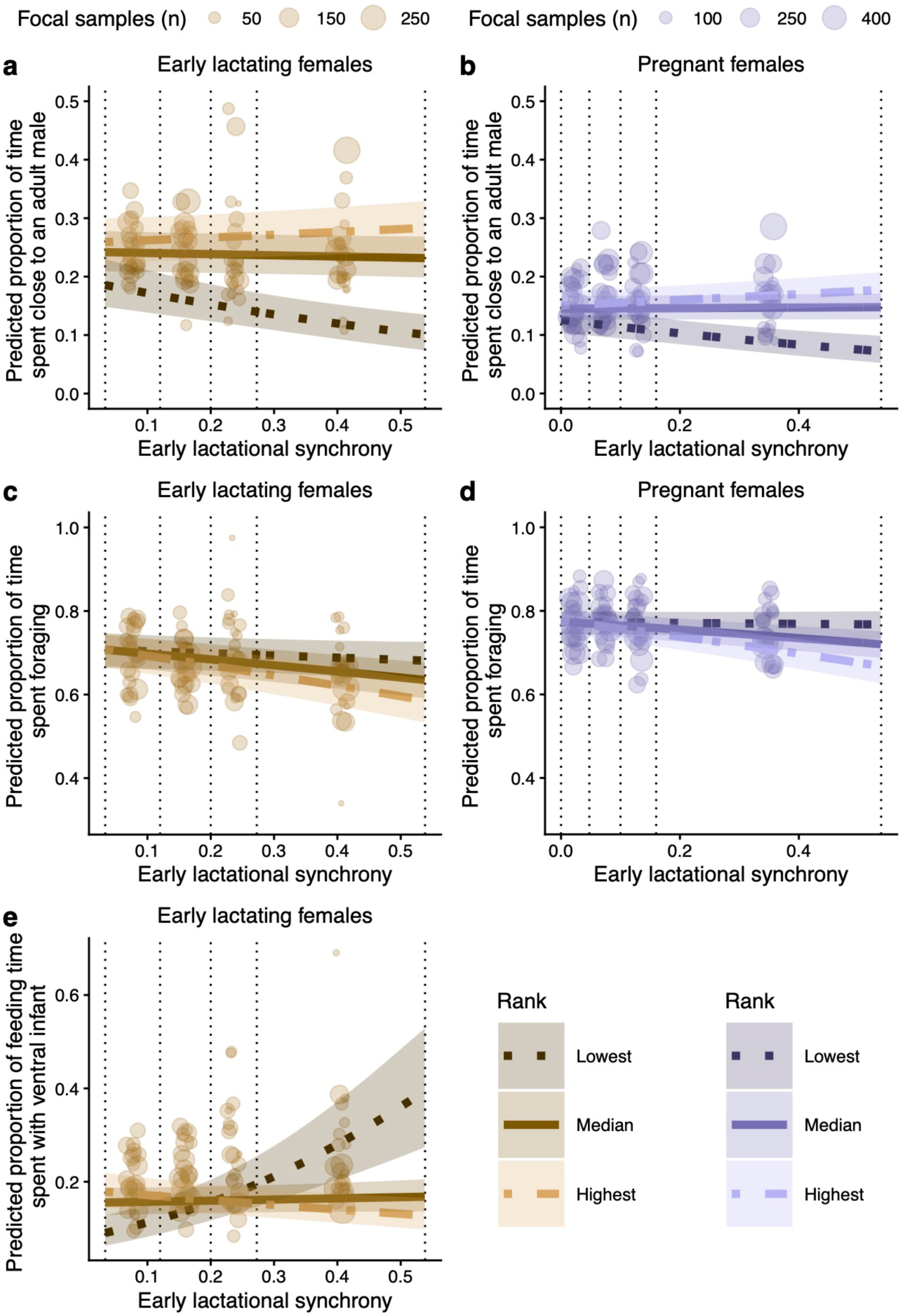
Rank-based differences in competition-related behaviors were more pronounced when early lactational synchrony was higher. (a-b) Higher-ranking females spent more time in proximity to adult males both in early lactation (a) and pregnancy (b), and this effect was strongest when early lactational synchrony was high. (c-d) High-ranking females spent less time foraging as early lactational synchrony increased, both during early lactation (c) and during pregnancy (d) but no such relationship existed for low-ranking females. (e) Low-ranking females in early lactation spent more of their feeding time in ventral contact with their infant when early lactational synchrony was higher, but high-ranking females did not. In all panels, the colored lines depict the GLMM-predicted relationship between early lactational synchrony and the response variable for females of the highest dominance rank (lightest-colored dot-dashed line; ordinal rank = 1 in panels a, b, and e; proportional rank = 1.0 in panels c and d), of the median rank observed in our dataset (solid line, ordinal rank = 7 in panels a, b, and e; proportional rank = 0.5 in panels c and d), and of the lowest dominance rank observed in our dataset (darkest-colored dotted line, ordinal rank = 29 in panels a, b, and e; proportional rank = 0.0 in panels c and d). The shaded regions around these lines represent the 95% confidence intervals of these predicted relationships. The colored circles represent the mean observed proportions calculated from all data collected on days that were within one quantile of early lactational synchrony in a complete hydrological year. These circles are proportionally scaled in size according to the number of focal samples contributing to the mean rate. The vertical dotted lines indicate the bounds of the quantiles of early lactational synchrony. For (a), and (c), n = 7,640 focal samples, for (b) and (d), n = 16,991 focal samples and for (e), n = 5,272 focal samples.

To explore the possibility that high early lactational synchrony also intensifies female-female competition over food, we examined the relationship between early lactational synchrony, female dominance rank, and time spent foraging (prediction 2.3). For both early lactating and pregnant females, we found that high-ranking females, but not low-ranking females, spent less time foraging as early lactational synchrony increased (Table 3; Fig. 4c,d; model 22, GLMM for early lactating females: proportional dominance rank: β = 0.030, 95% CI: [−0.161, 0.221]; early lactational synchrony: β = −0.241, 95% CI: [−0.561, 0.080]; interaction term: β = −0.819, 95% CI: [−1.356, −0.282]; model 23, GLMM for pregnant females: proportional dominance rank: β = 0.013, 95% CI: [−0.095, 0.120]; early lactational synchrony: β = −0.058, 95% CI: [−0.296, 0.180]; interaction term: β = −0.947, 95% CI: [−1.325, −0.568]). Neither rank nor early lactational synchrony were significant linear predictors of time spent foraging by early lactating or pregnant females, but their interaction term was well-bounded away from zero in both models. These foraging patterns may represent an indirect consequence of time spent with males: by spending more time with males in synchronous periods, high-ranking females may forage more efficiently and/or with fewer interruptions, thus reducing overall time spent foraging.

**Table 3.** Time spent foraging and feeding time spent in ventral contact with an infant are predicted by early lactational synchrony. Model-averaged estimates for the best-fitting models (ΔAIC_c_ < 2) predicting the proportion of time that early lactating females spent foraging (n = 7,640 focal samples of 235 early lactating females) and the proportion of feeding time that early lactating females spent in ventral contact with an infant (n = 5,272 focal samples of 234 early lactating females). The top 20 candidate models predicting these response variables can be found in the Supplementary Data file. Bold effects are those where the 95% CI does not overlap zero. Model-predicted values are presented to aid interpretation of estimates and were calculated for the extreme values of each significant fixed effect, while holding values for other fixed effects constant at their observed mean value.

| Response variable | Fixed effect | Estimate | Std.<br>Error | 95 % CI | Model-predicted values |
| --- | --- | --- | --- | --- | --- |
| Proportion of time spent foraging (model 22) | (intercept) | 0.834 | 0.131 | [0.578, 1.091] | — |
|  | Dominance rank | 0.030 | 0.097 | [-0.161, 0.221] | — |
|  | Early lactational synchrony | -0.241 | 0.163 | [-0.5661, 0.080] | — |
|  | **Dominance rank x Early lactational synchrony** | **-0.819** | **0.274** | **[-1.356, -0.282]** | Rank 0.0 (proportional, lowest), ELS 0.033 (min) = 0.707<br>Rank 0.0 (proportional, lowest), ELS 0.538 (max) = 0.681<br>Rank 1.0 (proportional, highest), ELS 0.033 (min) = 0.707<br>Rank 1.0 (proportional, highest), ELS 0.538 (max) = 0.586 |
|  | **Rainfall** | **-0.003** | **2.3 x 10<sup>-4</sup>** | **[-0.003, -0.002]** | 0.0 mm (min) = 0.696<br>286.7 mm (max) = 0.520 |
|  | Age | -8.0 x 10<sup>-4</sup> | 0.003 | [-0.006, 0.005] | — |
|  | Group size | 0.002 | 0.002 | [-0.003, 0.006] | — |
|  | Group size<sup>2</sup> | 9.4 x 10<sup>-6</sup> | 1.6 x 10<sup>-5</sup> | [-2.2 x 10<sup>-5</sup>, 4.1 x 10<sup>-5</sup>] | — |
| Proportion of feeding time spent in ventral contact with an infant (model 24) | (intercept) | -1.582 | 0.163 | [-1.902, -1.262] | — |
|  | Dominance rank | -0.033 | 0.008 | [-0.051, -0.016] | — |
|  | Early lactational synchrony | -0.947 | 0.324 | [-1.581, -0.312] | — |
|  | **Dominance rank x Early lactational synchrony** | **0.161** | **0.030** | **[0.102, 0.220]** | Rank 29 (ordinal, lowest), ELS 0.033 (min) = 0.090<br>Rank 29 (ordinal, lowest), ELS 0.538 (max) = 0.393<br>Rank 1 (ordinal, highest), ELS 0.033 (min) = 0.178<br>Rank 1 (ordinal, highest), ELS 0.538 (max) = 0.127 |
|  | **Proportion of sample with adult male** | **-0.305** | **0.057** | **[-0.416, -0.194]** | 0/10 min (min) = 0.169<br>10/10 min (max) = 0.131 |
|  | **Rainfall** | **0.007** | **4.6 x 10<sup>-4</sup>** | **[0.006, 0.008]** | 0.0 mm (min) = 0.139<br>268.7 mm (max) = 0.522 |
|  | Age | -0.001 | 0.003 | [-0.007, 0.006] | — |
|  | Group size | 0.001 | 0.002 | [-0.003, 0.004] | — |

As early lactational synchrony increased, early lactating females of low dominance rank exhibited strong increases in the proportion of feeding time that they spent in ventral contact with an infant, but no such pattern was apparent for mid- or high-ranking early lactating females (Table 3; Fig. 4e; prediction 2.4; model 24, GLMM; ordinal dominance rank: β = −0.033, 95% CI: [−0.051, −0.016]; early lactational synchrony: β = −0.947, 95% CI: [−1.581, −0.312]; interaction term: β = 0.161, 95% CI: [0.102, 0.220]). This observation may be related to the patterns we identified for time spent with adult males and time spent foraging: as low-ranking females are increasingly excluded from the protective benefits of male association, they may exhibit increasingly ‘restrictive’ maternal styles at a cost to their foraging efficiency. Specifically, ventral infant contact is likely most costly to foraging efficiency when feeding on grass corms (a critical fallback food that forms a large part of baboon diets during Amboseli’s long dry season^79–81^) and the lowest-ranking mothers substantially decreased their time spent feeding on grass corms when early lactational synchrony increased (see Supplementary Methods and Results; Fig. S4; models 28-31).

### High early lactational synchrony predicted increased infant mortality risk

Using data collected from 1976-2021 on 1,304 infants, we next employed a time-varying Cox proportional hazards model to test whether infant survival was compromised when early lactational synchrony was high (prediction 3.1; model 25). Controlling for other covariates thought to explain infant mortality, early lactational synchrony strongly predicted infant daily mortality risk during the first 90 days of life (Table 4; Fig. 5a; β = 3.011, 95% CI: [1.523, 4.500]). For instance, on days when a group was at our high synchrony benchmark, an infant’s mortality risk was predicted to be 1.9x higher than on days when the group was at the population median value of early lactational synchrony.

**Figure 5.**
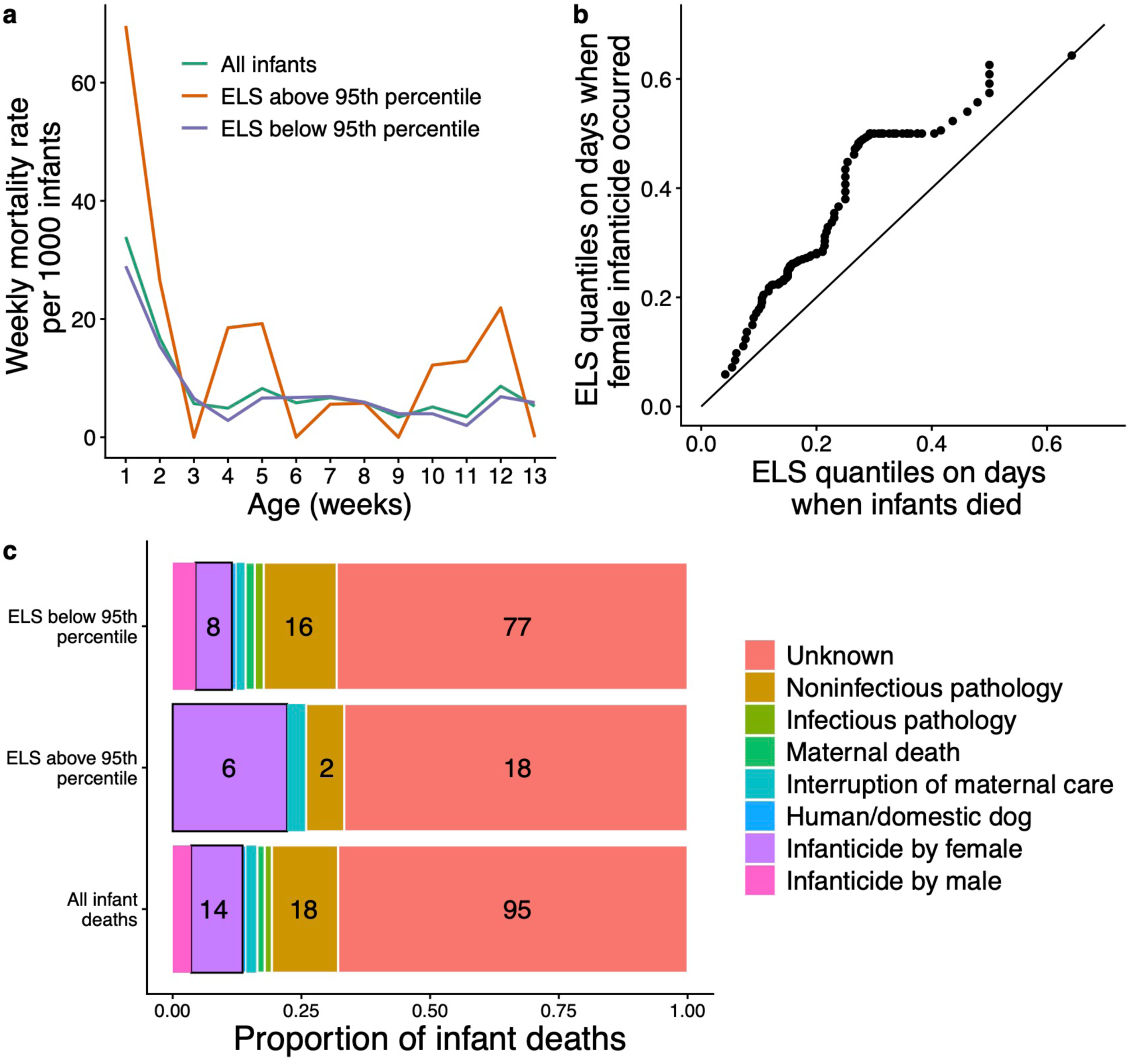
The relationship between early lactational synchrony (ELS), infant mortality, and infanticide. (a) Early lactational synchrony predicted infant mortality risk. Weekly mortality rates per 1000 infants are depicted with values calculated across all infants (green line), for infants in weeks in which the mean early lactational synchrony in their group was less than the 95^th^ percentile value of early lactational synchrony (purple line; <32.0% of females were in early lactation), and for infants in weeks in which the mean early lactational synchrony in their group exceeded the 95^th^ percentile value of early lactational synchrony (orange line; >32.0% of females were in early lactation). Early lactational synchrony was modeled as a continuous, daily, time-varying variable in our survival analysis; weekly mean values are used for visualization purposes. (b) Quantile-quantile plot showing that the distribution of early lactational synchrony is shifted higher on days when infants are killed by females (y-axis) than on days when infants die by any cause (x-axis). The solid, black line depicts the expected relationship if infanticide by females was equally likely on all infant death days. Almost all points fall above the line, indicating that days on which infants are killed by females tend to have higher early lactational synchrony relative to days on which infants die from any cause. (c) Assigned causes of infant death. Bar graphs depict the proportion of infant death causes for all infants included in the survival analysis (n = 140). The bottom bar includes all infants who died in their first 90 days of life during the study period (n = 140), the center bar only includes infants who died on days that were above the 95^th^ percentile value of early lactational synchrony (>32.0% of females in early lactation, n = 27), and the top bar only includes infants who died on days that were below the 95^th^ percentile value of early lactational synchrony (<32.0% of females in early lactation, n = 113). In all, infanticide deaths caused by females are highlighted with a black border. Numbers on the bar graphs indicate the sample sizes for the three most prevalent death causes (unknown, noninfectious pathology, and infanticide by females). Sample sizes for all death causes are given in Table S6.

**Table 4.** Infant mortality risk is predicted by early lactational synchrony. Model-averaged estimates for the best-fitting models (ΔAIC_c_ < 2) predicting the daily risk of all-cause infant mortality in the first 90 days after birth (model 25, Cox proportional hazards model, total infants = 1,304; died = 140, censored = 33; top 20 candidate models can be found in the Supplementary Data file). Effects where the 95% CI does not overlap zero are in bold. Recent cumulative rainfall was not retained in any of the best-fitting models. Model-predicted values are presented to aid interpretation of estimates and were calculated for the extreme observed values for each significant predictor variable for a 10-day-old infant, while holding values for other fixed effects constant at their observed mean value.

| Fixed effects | Estimate | Std.<br>Error | 95 % CI | Hazard<br>ratio* | Model-predicted values |
| --- | --- | --- | --- | --- | --- |
| **Maternal death** | **6.335** | **0.406** | **[5.539, 7.130]** | **563.691** | When an infant's mother is dead, mortality risk is 570.184 times higher than when she is alive |
| **Early lactational synchrony** | **3.011** | **0.759** | **[1.523, 4.500]** | **20.317** | At max. ELS (0.712), mortality risk is 7.702 times higher than at min. ELS (0.032) |
| Maternal age | -0.032 | 0.083 | [-0.194, 0.131] | 0.969 | — |
| Maternal age<sup>2</sup> | 0.001 | 0.003 | [-0.005, 0.008] | 1.001 | — |
| **Maternal rank** | **0.033** | **0.016** | **[0.003, 0.063]** | **1.034** | At lowest maternal rank (29), mortality risk is 2.562 times higher than at highest maternal rank (1) |
| Group size | -0.009 | 0.006 | [-0.021, 0.004] | 0.991 | — |
| Group size (time transformation term) | -0.003 | 0.008 | [-0.020, 0.012] | 0.996 | — |

\*A positive coefficient or a hazard ratio >1 means that higher values of the predictor (or the presence of the predictor in the case of maternal death) are associated with higher mortality hazard (lower infant survival), and negative coefficients or a hazard ratio <1 mean that higher values of the predictor are associated with lower mortality hazard (higher infant survival).

### Infanticide by females was a common source of infant mortality when early lactational synchrony was high

To understand the most important sources of infant mortality during periods of high early lactational synchrony, we reviewed the most probable causes of death for infants included in the survival analysis (i.e., those that died from 1976-2021). Deaths from infanticide were proportionally more common when early lactational synchrony was high (Fig. 5b,c; prediction 3.2; 22.2% of infant deaths on days above our high synchrony benchmark versus 11.5% of infant deaths on days below our high synchrony benchmark). Of the six infanticides known or strongly suspected to occur on days when groups exceeded our high synchrony benchmark, all six were known or very likely to be cases of infanticide by females living in the same group as the mother. When considering all infanticides committed by within-group females (13 total), 46.2% (six infanticides) occurred on days when groups exceeded our high synchrony benchmark. This proportion was significantly greater than the proportion of all infant deaths that occurred on days when groups exceeded our high synchrony benchmark (27/140 deaths, 19.3%; Fisher’s exact test: odds ratio = 3.549, *p* = 0.035). On the other hand, of the five total infanticides known or strongly suspected to be committed by males, none occurred on days when groups exceeded this threshold. Together, these patterns suggest that the relative concentration of infanticides in periods of high early lactational synchrony is explained by a rise in infanticidal behavior by females rather than males.

Notably, most cases of infanticide by females could not be attributed to direct resource competition among adult females at the time of the infant’s death. Rather, the majority of all within-group infanticides by females (11/13; 84.6%), including those performed during periods of high early lactational synchrony (4/6; 66.7%), were ‘kidnapping’ events performed by peripubertal, nulliparous females (prediction 3.3; Table S3). In these events, an immature female separated an infant from its mother for an extended period, and the infant subsequently died from starvation, dehydration, and/or rough handling. The remaining two within-group infanticides by females (15.4% of all known female infanticides and 33.3% of known female infanticides in periods of high early lactational synchrony) were thought to have occurred during episodes of severe, multiparty aggression that accompanied a permanent social group fission. Fully adult females were probably perpetrators in these cases, but their identities are unknown (Table S3).

## Discussion

These results provide evidence that group composition, specifically the proportion of adult females in early lactation, influences female-female competition and infant survival in baboons. Higher levels of early lactational synchrony were linked to more frequent agonistic interactions involving early lactating females. During periods of high early lactational synchrony, the costs and benefits of female social status were exaggerated, suggesting that early lactational synchrony increases female-female competition over access to protective male associates. High early lactational synchrony also strongly predicted infant mortality. Notably, infanticides, especially those performed by peripubertal females, were overrepresented in periods of high early lactational synchrony relative to infant deaths overall. While evidence of female-female competition for mates during the breeding season has been documented in several highly seasonally breeding mammal species^4,31–34^, our results provide strong evidence for negative fitness consequences of reproductive synchrony in a nonseasonal breeder. These fitness costs may select against strong breeding seasonality, even in highly seasonal environments.

Although it is seldom possible to definitively determine the cause of infant death, we suspect that synchrony-related infant mortality risk could be caused by several, co-acting phenomena. First, an increased risk of kidnapping by young, nonreproductive females during periods of high early lactational synchrony likely contributed to increased mortality overall. Second, these periods are also associated with increased rates of aggression involving mothers (Fig. 3, Table 2), potentially compromising the condition of dependent offspring. These two explanations are not mutually exclusive: aggressive contests involving mothers may also grant peripubertal females more opportunities to succeed in kidnapping attempts. Furthermore, while the energy required to produce milk and carry infants likely peaks after the 90-day window investigated here (because infants older than 90 days are both larger and more active)^42,43^, new motherhood nonetheless imposes high and rapidly increasing energetic demands and foraging challenges^41,82^. That females in early lactation allocated *less* time to foraging than females in any other reproductive state, potentially due to the constraints on time and physical mobility imposed by social vigilance and near-constant infant contact^41,46^, suggests that early lactation may be a particularly difficult time to meet increased energetic demands by increasing energy intake. Competition over food may thus arise when multiple females in the group face these challenges at the same time, potentially reducing maternal and offspring condition.

### Female agonistic behavior

Our finding that early lactating females were frequent participants in agonistic interactions (Figs. 2,3, Table 2) is consistent with observations in a large range of mammals that lactating females show heightened aggression relative to females in other reproductive states^83–85^. This phenomenon, known as ‘maternal aggression’ is understood as a mechanism to protect vulnerable offspring from conspecifics and predators. While maternal aggression may play a role in our study population, it cannot explain several of our results. Specifically, early lactating females did not initiate aggression more frequently in general than females in other reproductive states (see Supplementary Methods and Results, model 26), nor did they target their aggression toward the classes of females who posed the greatest threats to their infants (pregnant females and peripubertal females, models 12 and 27). Rather, they specifically targeted their aggression toward the class of females who shared a strong interest in maintaining proximity to adult males: other early lactating females (Fig. 2a).

While it has been proposed that female aggression directed toward yellow baboon mothers and their infants functions to reduce future competition^86,87^, we believe that competition over available male protectors is a more parsimonious explanation for the patterns of female aggression we observed, in line with the conclusions of work on chacma baboons^37,50^. Because protective male-female associations often form around the time of conception, peak in early lactation, and can persist until offspring independence^88^, the patterns of aggression we observed conform to expectations of competition for male social partners. Rank-based differences in time spent with males also provide indirect evidence of this competition. As Baniel et al.^50^ highlight, pregnant, lactating, and ovulating female baboons all potentially benefit from close association with males. Ovulating females may attract males more easily than pregnant or lactating females without the need to engage in much active competition. These dynamics may explain why pregnant females and early lactating females appear to compete with particular intensity for adult males (Fig. 2a,c and 4a-b, Table 2). Such an explanation avoids the need to invoke more complex cognitive mechanisms involving the projection of future competitive landscapes.

Notably, high early lactational synchrony not only predicted more frequent aggression in new mothers, but also strengthened the extent to which social status was associated with enhanced resource access (Fig. 4a,b). Adult males were monopolized by high-ranking females to a greater extent during periods of higher early lactational synchrony than otherwise. This effect may also have influenced female foraging patterns: when early lactational synchrony was higher, high-ranking females spent less time foraging and low-ranking females spent more of their feeding time in ventral contact with their infants (Fig. 4c,e). One explanation for this observation is that the benefits of male association (e.g., less frequent harassment^52^ or access to richer food patches^69^) were disproportionately allocated to high-status females. Those females may then have been freed to forage more efficiently than low-status females, who instead increased reliance on restricting infant spatial independence as a protective strategy. If restrictive mothering constrains food intake rates, this pattern could contribute to the extended interbirth intervals that low-ranking chacma baboons exhibit after giving birth in close temporal proximity to other mothers^37^. Synchrony-related restrictive mothering likely has the largest energetic impacts on low-ranking mothers during Amboseli’s long dry season, when the baboon population relies heavily on underground grass corms for food^79–81^. Ventral infant carrying likely imposes greater constraints on grass corm feeding than on any other type of feeding (Fig. S4a), and low-ranking mothers showed strong reductions in grass corm feeding time as early lactational synchrony increased (Fig. S4b).

Reduced foraging time by high-ranking females in periods of high early lactational synchrony could also result from increased investment in other behaviors (e.g., maintaining social relationships), or from increased monopolization of rich food patches while away from males. Alternatively, if reduced foraging time by high-ranking females reflects costs of maintaining association with males, rather than increased foraging efficiency, those costs may be outweighed, on balance, by the benefits of strengthening relationships with males during periods of intense female competition. Either way, we note that competition over current versus future resources and competition over food versus male associates are not mutually exclusive explanations for female agonistic and foraging behavior. Given that female competition for males likely impacts foraging behavior in baboons, it may be that resource competition and mate competition are more closely linked in female mammals than has been appreciated^5^.

### Infant survival and female infanticide

An unexpected and striking observation was the concentration of infanticidal behavior by peripubertal females within periods of high early lactational synchrony, despite the fact that both direct observations of infanticide and periods of high early lactational synchrony are rare in this population. Infant kidnapping and rough handling are well-known behaviors in female cercopithecine primates^89–91^, with some authors suggesting they may represent forms of reproductive competition^45,92^. Across social mammals, within-group infanticide by females is most common in groups that undergo periods of high birth synchrony, suggesting that it can result from competition over resources necessary for reproduction^7^. In this regard, our results are consistent with a general mammalian pattern of increased conspecific risk to infants when birth synchrony is high, and they provide a strong example of how group-level demographic conditions affect individual fitness.

However, while it is tempting to hypothesize that synchrony-related infanticidal behavior by peripubertal females is a tactic of direct competition with adult females, this hypothesis suffers from some logical flaws. If infanticide serves to reduce current competition, we would expect parous females rather than peripubertal (i.e., non-reproductive, nulliparous) females to commit infanticide when early lactational synchrony is high, especially because nulliparous females are unlikely to conceive in close temporal proximity to the births of the infants they target. Contrary to this expectation, parous females rarely engage in kidnapping, even when they have no current infants of their own. Furthermore, if infanticide serves to reduce the number of future competitors for the perpetrator or their offspring, neither peripubertal nor parous females should restrict their infanticidal attacks to periods of high early lactational synchrony, because the current number of young infants is unlikely to be a reliable predictor of future competition.

The drive to kidnap infants in peripubertal females is therefore unlikely to be directly related to competition among adult females. However, adult female competition could interact with the interest that peripubertal females commonly show in infants, if the frenetic nature of some aggressive interactions makes kidnapping attempts more likely to be successful. Furthermore, the relationship between infant kidnapping and early lactational synchrony may be compounded if some mother-infant pairs are competitively excluded from receiving male protection because the number of new mothers per male in the group is unusually high.

Whether early lactational synchrony itself increases interest in infants among peripubertal females remains an open question. It is possible that an unusual abundance of young infants may spur higher-than-usual interest in infants on the part of peripubertal females. If so, the probability that interactions with infants escalate to kidnapping may increase. An additional possibility is that, because infants of lower-ranking mothers are likely more easily kidnapped than infants of higher-ranking mothers, high early lactational synchrony may simply increase the probability that any given peripubertal female has access to infants of lower-ranking mothers. However, a *post hoc* analysis casts some doubt on this second possibility: during periods of high early lactational synchrony, an early lactating mother was *less* likely to have a higher-ranking peripubertal female as her nearest neighbor than during other periods (see Supplementary Methods; model 32, GLMM: β = −1.324, 95% CI: [−2.409, −0.240]). This pattern suggests that mothers avoid the most probable kidnappers in their group when early lactational synchrony is high. In support of this possibility (as opposed to general maintenance of distance from peripubertal females), early lactational synchrony did not predict whether early lactating mothers were likely to have *lower*-ranking peripubertal females as their nearest neighbors (model 33, GLMM: β = −0.141, 95% CI: [−0.822, 0.540]).

While infants born to low-ranking mothers were generally more likely to die than infants born to high-ranking mothers (Table 4), mothers of both high and low social status lost infants in periods of high early lactational synchrony (Fig. S5). This was somewhat surprising, given that low-ranking mothers appeared to bear more behavioral costs of high synchrony (Fig. 4a,c,e). Our investigation into infant death causes also suggested another surprising pattern: in the first three months of life, infants seemed to face a considerably higher risk of infanticide from female conspecifics than from male conspecifics (Table S3, Fig. 5c)^93^. We interpret this result with caution, given that the behavioral tactics of male vs. female infanticide may differ in ways that bias our probability of observing infanticidal acts (e.g., kidnappings tend to occur over several hours or days, while violent killings may be swift). However, we note this pattern in light of the historical emphasis on infanticide as a sexually-selected male behavior in research on social mammals^7,94,95^.

### Asynchronous breeding

Many mammals exhibit considerable plasticity in the extent to which reproduction is synchronized in response to seasonal variation in resource availability^36^, and comparisons between wild and captive breeding patterns indicate that primate breeding seasonality may be particularly flexible compared to some other mammalian orders^96^. A previous study on chacma baboons found that low-ranking females (but not high-ranking females) had longer interbirth intervals after periods of high reproductive synchrony, and that this rank-based difference could dampen breeding seasonality in an environment with high seasonal variance in resource availability^37^. In our study population, females exhibit stronger seasonal changes in the probabilities of experiencing menarche and postpartum cycling resumption than in the probabilities of conceiving and giving birth, which suggests that year-round, asynchronous breeding persists despite seasonal shifts in female energetic balance^75^. It remains to be seen whether the high costs of early lactational synchrony observed here explain this pattern. It also remains to be seen whether periods of high synchrony largely result from demographic stochasticity or from nonrandom alignment of female fertility within groups following extreme ecological or social events that correlate with fetal and/or infant loss^93,97^.

## Conclusions

The first detailed study of wild baboon maternal behavior concluded that “for the most part, maternity exaggerated the effects of dominance rank.”^41^ Our results extend this early finding from the individual level to the group level: baboon groups with more new mothers are characterized by dominance hierarchies that are more strongly predictive of females’ ability to access resources (even for non-mothers, Fig. 4b,d). They also highlight the important role that individual variation plays in producing the social structure of animal groups^40,98^, and they reveal that emergent social structure can have strong effects on individual fitness. As such, the potential for demographic variability to affect competitive regimes and the potential for subsequent group-level variation in competitive regimes to affect individual fitness outcomes should be investigated in a broader range of group-living species. Among social mammals, within-group female-female competition and female infanticide are widespread^7^, despite the prevalence of female philopatry and female-biased kinship^99,100^. Therefore, species in which philopatric females engage in reproductive competition with behavioral tactics up to and including infanticide^7,101^ are of particular interest. We suggest that future studies should focus on the proximate factors shaping female aggression, the underlying mechanisms motivating juvenile and adult female interactions with non-offspring young, and the roles of inter- and intrasexual social interactions in shaping the timing of reproduction.

## Methods

### Study site and subjects

This population of wild baboons lives in the Amboseli basin, a semi-arid, short-grass savannah ecosystem at the northern base of Mt. Kilimanjaro in southern Kenya. The basin experiences an annual dry season from June to October when nearly no rain falls, followed by a highly variable season from November to May when the amount of rainfall varies greatly from month to month and year to year^79^. The baboon population has been continuously monitored by the Amboseli Baboon Research Project (ABRP) since 1971^72^. Over the course of the study’s history, the number of habituated social groups monitored at any one time has ranged from one to six, and the groups have undergone multiple fissions and fusions since the late 1980s^81^. See Supplementary Methods for additional details on daily demographic, behavioral, and ecological monitoring by the ABRP.

### Female reproductive states and lactational synchrony

In our analyses, we considered both individual female reproductive states and a group-level metric of reproductive state similarity as important predictors of individual behavior. We identified three ‘major’ female reproductive states (ovarian cycling, pregnant, and postpartum amenorrhea^102^), each of which can be subdivided further into finer-grained ‘minor’ reproductive states (Fig. 1; see also Supplementary Methods). Throughout, we use the term early lactational synchrony to refer to the proportion of adult females in a group that were in early lactation, i.e., had an infant <90 days old. Further information on how early lactational synchrony was calculated can be found in the Supplementary Methods.

Our analyses of both competitive behavior and infant survival rely on the assumption that the 90-day period after birth is particularly relevant to female-female competition. To further interrogate this assumption beyond our primary analyses of competition-related behavior (see sub-hypothesis 1), we constructed several alternative statistical models for the analyses predicting agonistic interaction rates and infant survival (models 34-39). In each of these models, we replaced the early lactational synchrony fixed effect with a term for group-level synchrony in one of the other ‘minor’ reproductive states included in postpartum amenorrhea (early-mid lactation, late-mid lactation, and late lactation). These were calculated analogously to our calculation of early lactational synchrony (e.g., the proportion of adult females in a group that were in early-mid lactation on a given day). We found no evidence that synchrony in any other stage of postpartum amenorrhea besides early lactation was associated with either increased agonistic behavior by new mothers or increased infant mortality risk (although higher synchrony in early-mid lactation was significantly associated with *decreased* agonistic behavior by new mothers; see model 34).

### Competition-related behaviors

To test whether early lactation is a particularly competitive time for adult females, we calculated four competition-related metrics from data collected during 10-minute focal animal samples from October 1999 to December 2021, for females in each of the 12 ‘minor’ reproductive states (see Supplementary Methods for more details on focal sampling). These were the: (i) rate of agonistic interactions, (ii) proportion of observed feeding time spent in ventral contact with an infant, (iii) proportion of observation time spent foraging, and (iv) proportion of observation time spent within 5 m of at least one adult male. To test probable drivers of competition among females, we modeled these four variables as functions of female reproductive state, early lactational synchrony, and other covariates (see below for details on statistical modelling). For each focal sample, we calculated agonistic interaction rate by summing the number of intragroup dyadic agonistic interactions occurring between a focal female and a nonfocal adult female and dividing it by the number of minutes in the focal sample. Agonistic interactions were only included in this rate if either one female displayed aggressive behaviors and the other displayed submissive behaviors or if one female spatially displaced the other. Foraging time was defined as the sum of time spent feeding and time spent walking while not feeding, as in^79,103,104^. Male neighbors were considered adults once they attained a higher dominance rank than at least one other adult male^105^. Sample sizes for each of the 12 ‘minor’ reproductive states are given in Table S4.

### Other predictors of competition-related behavior and infant survival

In addition to reproductive state, we considered dominance rank and age as likely individual-level predictors of competition-related behavior in females. Female baboons form linear dominance hierarchies in which higher ranking individuals have priority of access to resources, including food and male social partners^51,106^. In all statistical models (see below), we used Akaike’s information criterion (AIC_c_) values to compare candidate models with female dominance rank coded as either a proportional rank or ordinal rank, as previous research has demonstrated that many behavioral metrics are better predicted by either one rank metric or the other^107^. In all cases, we report the results for the model with the lower AIC_c_ value. Consistent with previous findings^107^, proportional rank was a better predictor of foraging behavior and rates of female-female agonistic interactions, while ordinal rank was a better predictor of time spent with males, feeding time spent in ventral contact with an infant, and infant survival. Because female social connectedness declines with age in this population^51^, we expected older females may engage in fewer social interactions overall than younger females (including agonistic interactions and socio-spatial proximity). We expected infant mortality risk would be higher both for younger, inexperienced mothers and older, senescent mothers.

For each group on each day, we calculated the number of adult females in each reproductive state and the number of adult males to control for between-focal sample variation in the opportunity for focal females to socially interact with specific classes of individuals. We also expected that total group size (after controlling for the number of potential interactors of a given sex or state class) might affect some of our behavioral variables of interest. Specifically, foraging-related behaviors have been shown to vary quadratically with group size in this population^103,108^. We also expected that, after controlling for the number of potential agonistic partners, rates of agonistic interaction might vary nonlinearly with total group size. Specifically, we expected that rates of agonistic behavior involving focal females (per nonfocal female) would initially decrease with increasing group size, because time budget constraints and/or the increasing costs of injury risk would prevent individual females from additively increasing investment in aggression in larger groups. Based on preliminary data visualizations (Fig. S6), we thought this trend might level off or even reverse as groups became very large, perhaps due to high within-group competition. Finally, we expected that the proportion of feeding time that females spent in ventral contact with an infant would increase with group size, as the total number of potential conspecific threats to the infant became higher.

Baboons rely on a broad range of diet items which can vary dramatically in their availability throughout the hydrological year and in the extent to which their ‘patches’ can be monopolized by dominant individuals^79,106^. Therefore, it is challenging to make simple predictions about the relationships between the availability of specific food types in the environment and the probability of feeding-related aggression. For our analyses, we considered recent cumulative rainfall as a rough approximation of general food availability. To determine an appropriate time window over which to measure cumulative rainfall, we calculated the correlation coefficient between the proportion of time spent foraging in a focal sample (with the assumption that individuals would spend more time foraging when food availability and/or quality was lower) and four potential windows of recent cumulative rainfall (30, 60, 90, and 120 days prior to the focal sample). Cumulative rainfall in the 30 days prior to a focal sample had the strongest negative correlation with time spent foraging, and this correlation became weaker when measuring cumulative rainfall over longer windows.

### Statistical approach

We took an information theoretic approach to all analyses, which were conducted in R v4.5.0^109^. We began by constructing ‘full models’ that included all likely covariates. We then used the *dredge* function in the *MuMIn* R package^110^ to acquire AIC_c_ values for all models with all possible combinations of fixed effects included in the full model^111,112^. All candidate models within 2 AIC_c_ values of the candidate model with the lowest AIC_c_ value were selected as the best-fitting models. We used *MuMIn*’s *model.avg* function to obtain a weighted average of the best-fitting models (those within 2 AIC_c_ of the best model) and estimate effect sizes for covariates. If a fixed effect was left out of a particular candidate model, then a weighted effect size of zero was included in the calculation of the overall weight-averaged effect size for that fixed effect. We considered fixed effects to be statistically significant when the 95% confidence interval of their weight-averaged effect size did not overlap zero. Cox proportional hazards models were constructed using the *coxph* function from the package *survival*^113^ and GLMMs were constructed using the package *glmmTMB*^114^.

### Sub-hypothesis 1: characterizing competition-related behavior across the interbirth interval

To test our prediction that females in early lactation experience agonistic interactions more frequently than females in other reproductive states (a proxy of direct competition, Prediction 1.1), we constructed a negative binomial GLMM with a log-link function predicting the number of agonistic interactions with other adult females that focal females experienced during a focal sample (model 1, n = 60,655 focal samples). We chose to model these count data with a negative binomial distribution because they were overdispersed. In the model we included the following fixed effects: (i) the focal female’s ‘minor’ reproductive state, her proportional dominance rank as a (ii) linear and (iii) quadratic term (because we expected both high-ranking and low-ranking females would be involved in more agonistic interactions than mid-ranking females), (iv) her age, (v) the number of adult females (i.e., potential interactors) in her group, the number of total individuals her in group as a (vi) linear and (vii) quadratic term (because of the potential nonlinear effects of total group size, after controlling for the number of potential interactors, Fig. S6), and (viii) the cumulative amount of rainfall that had fallen in the previous 30 days. We also included the log-transformed number of minutes that the focal female was in sight of the observer during the focal sample as an offset term. We included focal female identity, group identity, and hydrological year as random intercepts to control for repeated sampling of females and groups, and for potential variation between females, groups, and years that were not captured by our fixed effects. Because no female-female agonistic interactions occurred in the vast majority of 10-min focal samples, we used the *check_zeroinflation* function in the R package *performance*^115^ to test whether the model was significantly zero-inflated. However, no zero-inflation was detected (observed number of zeros = 57,920; predicted number of zeros = 57,570; ratio = 1.00; *p* = 0.672).

To test Prediction 1.2, we constructed a binomial GLMM with a logit-link function predicting the proportion of observed feeding time that a focal female spent in ventral contract with an infant during a focal sample (model 2, n = 45,706 focal samples in which any feeding was observed). Fixed effects included: (i) the focal female’s ‘minor’ reproductive state, (ii) her ordinal dominance rank, (iii) her age, (iv) the proportion of all minutes in the sample that she spent within 5 m of an adult male (because infant spatial independence increases in the presence of adult males^116^), (v) the total size of her group, and (vi) the cumulative amount of rainfall that had fallen over the previous 30 days. Again, we included focal female identity, group identity, and hydrological year as random effects. Because some candidate models constructed from these covariates failed to converge, we rescaled numeric fixed effects and centered them at zero, which remedied convergence issues.

To test Prediction 1.3, we constructed a binomial GLMM with a logit-link function predicting the proportion of observation time that focal females spent foraging during a focal sample (model 3, n = 60,665 focal samples). Here, we included (i) the focal female’s ‘minor’ reproductive state, (ii) her proportional dominance rank, (iii) her age, the total size of her group as a (iv) linear and (v) quadratic term, and (vi) the cumulative amount of rainfall that had fallen over the previous 30 days as fixed effects and focal female identity, group identity, and hydrological year as random effects.

Similarly, to test Prediction 1.4, we constructed a binomial GLMM with a logit-link function predicting the proportion of observation time that focal females spent within 5 m of an adult male (model 4, n = 60,655 focal samples). Fixed effects in this model were (i) the focal female’s ‘minor’ reproductive state, (ii) her ordinal dominance rank, (iii) her age, (iv) the number of adult males (i.e., potential neighbors) present, (v) the number of nonfocal cycling females present (because they are attractive alternative proximity partners for potential male neighbors), and (vi) the cumulative amount of rainfall that had fallen over the previous 30 days. Random effects included focal female identity, group identity, and hydrological year.

Different explanations for female-female aggression (e.g., competition over food vs. mates, current competition vs. future competition reduction) offer different predictions about how females in different reproductive states should participate in agonistic interactions^47^. Therefore, we also modeled rates of agonistic interactions initiated toward and received from females in each of four mutually exclusive reproductive states: cycling, pregnant, early lactation, and the entire period of postpartum amenorrhea following early lactation (eight separate models, models 5-12). Each of these eight models was fit using a negative binomial GLMM with a log-link function predicting the number of times that a focal adult female (of any reproductive state) either initiated an agonistic interaction toward or received an agonistic interaction from a female in one of these four states. In each case, we included the following fixed effects: (i) the focal female’s proportional dominance rank, (ii) her own reproductive state, (iii) her age, (iv) the number of nonfocal females who were in the reproductive state of interest, the total group size as a (v) linear and (vi) quadratic term, and (vii) the cumulative amount of rainfall that had fallen in the previous 30 days. As in our model predicting total agonistic interactions, we included an offset term for the log-transformed number of minutes that the focal female was in sight of the observer during the focal sample and random effects for focal female identity, group identity, and hydrological year.

In these models, we only included focal samples in which at least one nonfocal female of the reproductive state of interest was present in the group, and thus could engage in agonistic interactions with the focal female. Therefore, sample sizes differ across these models (n = 48,335 focal samples for models predicting interaction rates with early lactating females; n = 56,635 focal samples for models predicting interaction rates with post-early lactation postpartum amenorrhea females; n = 59,679 focal samples for models predicting interaction rates with cycling females; n = 58,602 focal samples for models predicting interaction rates with pregnant females). In some cases, some candidate models predicting reproductive state-specific agonistic interaction rates failed to converge, likely because of the model structure complexity given the relative infrequency of events. See Supplementary Methods for further information on how we handled these cases.

### Sub-hypothesis 2: early lactational synchrony and female-female competition

To test whether early lactational synchrony increases female-female competition (sub-hypothesis 2), we modeled our four competition-related behavioral metrics (agonistic interaction rates, feeding time spent in ventral contact with an infant, time spent foraging, and time spent with adult males) as functions of early lactational synchrony. To test prediction 2.1, we constructed a negative binomial GLMM with a log-link function in which the response variable was the number of agonistic interactions that a focal early lactating female initiated with other adult females during a focal sample (model 13, n = 7,640 focal samples). We included the same fixed and random effects and offset term that were in the models predicting agonistic interactions initiated toward females in specific reproductive states (models 5-12), but with an additional fixed effect of (viii) the proportion of adult females in the group who were in early lactation. We also substituted the number of all nonfocal adult females present for the number of nonfocal females in the specific state of interest.

After determining that early lactating females initiated more agonistic interactions when early lactational synchrony was high, we sought to determine whether this increase was solely driven by an increase in the total number of individual targets (i.e., other early lactating females; Fig. 2a), or if early lactating females also increased rates of agonism with individual females of particular reproductive states in periods of high synchrony. To do so, we constructed four negative binomial GLMMs predicting the number of agonistic interactions that early lactating females initiated toward nonfocal early lactating females (model 14), cycling females (model 15), pregnant females (model 16), and post-early lactation postpartum amenorrhea females (model 17). These models included the same fixed and random effects and offset term as in model 13, but controlled for the number of nonfocal females in the reproductive state of interest, rather than the number of all nonfocal adult females. Again, we only included focal samples if a nonfocal female of the reproductive state of interest was present (n = 6,248 focal samples for interactions with nonfocal early lactating females; n = 7,504 focal samples for interactions with post-early lactation postpartum amenorrhea females; n = 7,444 focal samples for interactions with pregnant females; n = 7,536 focal samples for interactions with cycling females). In cases where some candidate models failed to converge, we followed the same procedure as for the models predicting general rates of agonistic interactions with females of specific reproductive states (see Supplementary Methods for details).

Next, we tested if the rise in female-female aggression with early lactational synchrony could be related to competition over male social partners or food (predictions 2.2-2.3). To do so, we constructed five binomial GLMMs with logit-link functions, each of which tested the relationship between early lactational synchrony and either the proportion of observation time that females spent within 5 m of an adult male, the proportion of observation time that females spent foraging, or the proportion of observed feeding time that females spent in ventral contact with an infant. We tested the foraging time response variable and the male proximity time response variable with two models each: one including data from focal samples conducted on females in early lactation (n = 7,640 focal samples on 235 females) and one including data from focal samples conducted on pregnant females (n = 16,991 focal samples on 250 females). We examined behavior during these two reproductive states because females in these states initiated agonistic interactions with females in early lactation most frequently (Fig. 2a). Because pregnant females never had young infants to carry ventrally, we only used data from focal samples on early lactating females to test the relationship between early lactational synchrony and feeding time spent in ventral contact with an infant. We also only used focal samples in which the focal early lactating female fed (n = 5,272 focal samples on 234 females).

For the two models predicting time spent in close proximity to males (prediction 2.2, models 18-19), we included seven fixed effects: (i) the proportion of adult females in the group who were in early lactation, (ii) the focal female’s ordinal dominance rank, (iii) an interaction term between (i) and (ii), (iv) the focal female’s age, (v) the number of adult males present, (vi) the number of cycling females present, and (vii) the cumulative amount of rainfall that had fallen over the last 30 days. We modeled an interaction between early lactational synchrony and dominance rank to test whether dominance rank-related differences in time spent with adult males would become stronger when early lactational synchrony was higher, a prediction based on the hypothesis that early lactational synchrony intensifies competition over access to males (prediction 2.2). In both models, we included focal female identity, group identity, and hydrological year as random effects. We also investigated the possibility of an additional interaction effect between the number of adult males in the group and early lactational synchrony on the proportion of time spent with males. All statistical results of this analysis are given in the Supplementary Data file (models 20-21) and described in Fig. S7.

For the two models predicting time spent foraging (prediction 2.3, models 22-23), we included seven fixed effects: (i) the proportion of adult females in the group who were in early lactation, (ii) the focal female’s proportional dominance rank, (iii) an interaction term between (i) and (ii), (iv) the focal female’s age, the total group size as a (v) linear and (vi) quadratic term, and (vii) the cumulative amount of rainfall that had fallen in the last 30 days. Focal female identity, group identity, and hydrological year were included as random effects.

For the model predicting feeding time spent with a ventral infant (prediction 2.4, model 24), we included seven fixed effects: (i) the proportion of adult females in the group who were in early lactation, (ii) the focal female’s ordinal dominance rank, (iii) an interaction term between (i) and (ii), (iv) the focal female’s age, (v) the proportion of all sample minutes in which the focal female was within 5 m of an adult male, (vi) total group size, and (vii) the cumulative amount of rainfall that had fallen over the last 30 days. Again, random effects included were focal female identity, group identity, and hydrological year.

### Sub-hypothesis 3: infant survival and infanticide risk

To test our prediction that high early lactational synchrony is associated with an increased risk of infant death, we used a Cox proportional hazards model with time-varying covariates to predict the daily risk of infant death in the first 90 days of life (model 25). Our covariates were: (i) the proportion of adult females in early lactation on each day of the infant’s life; (ii) group size; (iii) whether or not the infant’s mother died on or before that day; (iv) maternal ordinal rank; maternal age as a (v) linear and (vi) quadratic term; and the (vii) cumulative total rainfall that fell in the last 30 days. While the terms for early lactational synchrony, group size, maternal death, maternal rank, and cumulative rainfall were allowed to vary each day of an infant’s life, maternal age and maternal rank were treated as fixed variables and were coded as the value corresponding to the infant’s birthdate. Infant identity was included as a cluster term. To test whether any covariates violated the assumption of proportional hazards, we used the *cox.zph* function of the R package *survival*^113^ to test for a non-zero slope in the scaled Schoenfeld residuals. Non-proportionality was detected for the covariates maternal age (*p* = 0.006), maternal age^2^ (*p* = 0.010), group size (*p* = 0.006), and maternal death (*p* = 0.003). Therefore, we applied a time transformation on each of these covariates in our full survival model, which allows their estimated effect sizes to vary with infant age, instead of assuming a constant effect across infant development. Because not all time transformation variables were retained in the ‘best’ survival models, we present the full survival model with all time transformations in Table S5. The effect of our variable of interest (early lactational synchrony) is statistically significant and of a similar magnitude in both the full model (Table S5) and in the weighted average of the best-fitting candidate models (Table 4). In total, 1,304 infants born between November 1976 and December 2021 were included in this analysis, of whom 140 died in the first 90 days of life. An additional 30 infants were still alive at their last observation but were treated as right-censored because they could not be followed through the full 90-day period. We restricted our survival analyses to this period because rainfall data were not recorded before November 1976.

After observing a negative relationship between early lactational synchrony and infant survival, we sought to investigate the causes of death for infants who died during periods of high early lactational synchrony (see Supplementary Methods for information on how we assigned causes of death and confidence in the assigned causes). Of the 140 infant deaths included in our survival analysis, causes of death with our highest confidence level could be assigned to 45 individuals.

Following this assignment process, we visualized the relative proportion of each cause of death for (i) all infants, (ii) those infants who died on days that were below the 95^th^ percentile value of early lactational synchrony (<32.0% of females in the group in early lactation, n = 113 deaths), and (iii) those infants who died on days that were above 95^th^ percentile value of early lactational synchrony (>32.0% of females in the group in early lactation, n = 27 deaths, Fig. 5c). In this assessment, any infant whose cause of death was not assigned with our highest level of confidence was considered to have died of an unknown cause. Infanticide deaths were overrepresented among deaths on days that were above the 95^th^ percentile value of early lactational synchrony, both relative to all deaths and relative to deaths on days below the 95^th^ percentile value. This observation led us to review field notes related to each infanticide death to determine the most likely actors (male or female) and contexts (within-group or between-group) of infanticide when early lactational synchrony was high (Table S3). In assigning causes of death, we use the definition of infanticide proposed by Digby^117^ and more recently invoked by Lukas and Huchard^7^: ‘an act that makes a direct or significant contribution to the immediate or imminent death of conspecific young.’ This definition includes acts of physical aggression toward infants and acts such as kidnappings that cause infants to die via ‘enforced neglect’^7^. It thus focuses on the ultimate consequences of interactions with conspecific young, regardless of proximate motivation.

### Animal ethics

Data collection protocols used for this study were approved by the Institutional Animal Care and Use Committees (IACUC) at Duke University, the University of Notre Dame, Princeton University, and the Ethics Council of the Max Planck Society and adhered to all the laws and guidelines of Kenya. All data collection for this study was observational, and study subjects showed no signs of distress associated with observation.

## Data Availability

Data necessary to reproduce these analyses are currently available for reviewers on the Dryad Data Repository under DOI:10.5061/dryad.83bk3jb26^118^. These data and code will be made publicly available at Dryad Data Repository upon publication.

## Code Availability

*R* code necessary to reproduce these analyses are available alongside the published Dryad data set^118^, and at Zenodo under DOI:10.5281/zenodo.13737391^119^.

## Acknowledgments

We thank Jeanne Altmann for her pivotal role in stewarding the Amboseli Baboon Research Project and in designing many of the ABRP’s long-term data collection protocols. We thank the Max Planck Institute for Evolutionary Anthropology, Duke University, Princeton University, the University of Notre Dame, and Stony Brook University for financial and logistical support. In Kenya, our research was approved by the Wildlife Research Training Institute (WRTI), Kenya Wildlife Service (KWS), the National Commission for Science, Technology, and Innovation (NACOSTI), and the National Environment Management Authority (NEMA). We also thank the University of Nairobi, the Kenya Institute of Primate Research (KIPRE), the National Museums of Kenya, the members of the Amboseli-Longido pastoralist communities, the Enduimet Wildlife Management Area, Ker & Downey Safaris, AirKenya, and Safarilink for their cooperation and assistance in the field. We are particularly grateful to Raphael Mututua, Serah Sayialel, and Lilian Musembei for their work as members of the ABRP long-term field team, to Tim Wango and Vivian Oudu for assistance in Nairobi, and to Jacob Gordon, William Wilbur, and Caitlin Broderick for management of the baboon project database, BABASE. Database design and programming were provided by Karl Pinc. We also thank Andreas Koenig, Heather Lynch, Leonie Pethig, Claudia Fichtel and four anonymous reviewers who provided feedback on earlier versions of this manuscript, as well as the Behavioral Ecology Group at Stony Brook University for discussion at an early stage of this analysis.

