## Supplementary Information for "High early lactational synchrony within baboon groups predicts increased female-female competition and infant mortality"

### Supplementary Methods

*Daily demographic, behavioral, and ecological monitoring*. The Amboseli Baboon Research Project monitored between one to six social groups at any one time over the course of the study (begun in 1971; 1976-2021 are included in this data set). The number of groups monitored has varied over the years, and therefore the frequency with which any given study group was observed ranged from 2-4 days per week to near-daily. The typical inter-observation gap was 1-3 days. During each visit, researchers conducted a census of the group members present, recorded notes on any demographic events (e.g., births and deaths), and collected reproductive data on each female. Infant birth and death dates were assigned as the date on which the birth or death occurred. In the case of births and deaths that happened during observational gaps, dates were assigned based on the most likely date given the available clues or, lacking clues, as the midpoint between the last observation date prior to birth or death and the first observation date after it. Female reproductive state (cycling, pregnant, or postpartum amenorrhea) was inferred for each day from information on paracallosal skin color, presence of visible menstrual blood, sexual swelling size, and the presence of a suckling infant. The reliability of inferring female reproductive state from these characteristics has been confirmed by hormonal analyses[1-3]. Reproductive and behavioral data were sparse for some time periods, particularly during permanent group fission events, when groups are often split into subgroups and not all individuals can be located. Therefore, we excluded these periods from our analyses.

From October 1999 to December 2021, observers conducted 10-minute focal follows of all adult females in a randomized order throughout the observation day. During focal samples, researchers recorded female activity states, postural position (sitting vs. standing), near neighbor information, and, if applicable, infant contact status (ventral vs. dorsal) at 60-second intervals. Near neighbor information included: (i) the identity of the focal female's nearest neighbor (of any age-sex class) within 5 m, (ii) the identity of the focal female's nearest adult neighbor within 5 m, and (iii) the identity of the focal female's nearest adult neighbor within 5 m who was of the opposite sex of the adult neighbor identified in (ii). Additionally, researchers collected data on all observed agonistic interactions involving the focal female, including information on the identities of participants and any aggressive or submissive behaviors displayed. All adult females were assigned dominance ranks according to a monthly dominance matrix based on wins and losses in female-female agonistic interactions[4,5]. Rank assignments were made using agonistic behavior data recorded both in and out of the context of focal animal samples, but calculations of rates of agonistic behavior were made exclusively with data collected during focal sampling. Because focal samples were collected on all adult females according to a randomized order, sampling effort was evenly distributed across high-, mid-, and low-ranking females (see Fig. S7 for the distribution of ordinal and proportional dominance of focal females included in our analyses). Since 1976, researchers have also recorded the daily rainfall in millimeters using a rain gauge located in the field camp.

*Defining reproductive states.* In our analyses, we categorize female reproductive states into three 'major' categories, each of which can be subdivided into finer-grained 'minor' categories:

(i) Ovarian cycling lasted from the first day a female's sex skin was turgescent after menarche or the end of her last pregnancy to either the next onset of pregnancy or death. Subdivisions of ovarian cycling include 'flat' (days when females do not exhibit any sexual swelling), 'turgescence-follicular' (lasting from the first day a female's sex skin was turgescent to the sixth day before her sex skin became deturgescent), 'turgescence-ovulation' (lasting from the fifth day before a female's sex skin became deturgescent to the day before her sex skin became deturgescent), and 'deturgescence-luteal' (lasting from the first day a female's sex skin was deturgescent to the last day any swelling was discernible)[3].

(ii) Pregnancy lasted from the first day a female's sex skin was deturgescent in her last sexual cycle preceding a known pregnancy to the day before she gave birth, experienced fetal loss, or died. Pregnancy was subdivided into three 60-day trimesters (mean ± SD duration of pregnancy: 178 ± 6 days[2]; first trimester: day 1-60; second trimester: day 61-120; third trimester: day 121-parturition).

(iii) Postpartum amenorrhea lasted from the day a female gave birth or experienced fetal loss to the next day her sex skin was turgescent or she died. The duration of postpartum amenorrhea is far more variable than the duration of pregnancy (mean ± SD duration of postpartum amenorrhea: 322 ± 87 days[2]). For the first 360 days of postpartum amenorrhea, we defined 90-day subdivisions of equal length ('early lactation': day 1-90; 'early-mid lactation': day 91-180; 'late-mid lactation': day 181-270; 'late lactation': day 271-360). We pooled all females who had not yet resumed cycling more than 360 days after giving birth into a single subdivision: 'post-year postpartum amenorrhea'. While milk production may continue beyond 360 days after birth, Altmann[6] estimated that yearling baboons were, on average, capable of intaking adequate amounts of energy and protein from independent foraging by this age. Altmann[6] provides 35 weeks as a rough estimate of peak milk volume production by baboon mothers in Amboseli, corresponding to our 'late-mid lactation' category.

*Quantifying early lactational synchrony*. For each study group on each day of its inclusion in this study, we calculated early lactational synchrony as the proportion of all adult females in the group who were in the first 90 days of postpartum amenorrhea with a live infant. Early lactational synchrony was only calculated on a given day if the reproductive state of all adult (i.e., post-menarche) females in the group was known and the group was not known to have split into subgroups. We were able to gather this information for 58,322 group-days (159.68 group-years) between 1976 and 2021 across a total of 19 distinct social groups (two original study groups and their descendant groups, which persisted for varying lengths of time), for an average of 3,069.58 days (8.40 years) per group. Note that because early lactational synchrony is calculated as the proportion of adult females in a group that were in early lactation, some values are inherently unlikely due to the distribution of adult female group sizes in this population (e.g., a value of exactly 0.01 would necessitate a group size containing 100 adult females, far more than we ever observe).

We based our 90-day definition of early lactation on careful consideration of the large body of published research on baboon maternal behavior and infant development. However, to confirm that 90 days postpartum corresponds to the timing of key changes in competition-related behavior for baboon mothers, we used our data set of 24,671 focal animal samples of 237 females in postpartum amenorrhea to visualize changes in our four behaviors of interest (agonistic interaction rate, proportion of feeding time spend in ventral contact with an infant, proportion of time spent foraging, and proportion of time spent within 5 m of an adult male) over the first 360 days of postpartum amenorrhea. For every focal sample conducted on a female in the first 360 days of postpartum amenorrhea, we identified the number of days that had passed since the focal female most recently gave birth. We compiled every focal sample collected on each unique ordinal day from day #1 to day #360 of postpartum amenorrhea (mean = 63.928 ±

21.084 s.d. samples per day) and calculated the mean agonistic interaction rate, mean proportion of feeding time spent in ventral contact with an infant, mean proportion of time spent foraging, and mean proportion of time spent within 5 m of an adult male on each day since parturition across those samples. Next, we compiled focal samples into 10-day bins (day #1 to day #10; day #11 to day #20; etc.) and calculated these same metrics over each 10-day interval (mean = 639.278 ± 198.590 s.d. focal samples per 10-day bin). We plotted these values against time since parturition (Fig. S1-2) and visually assessed how changes in each behavior of interest aligned with our expected 90-day threshold.

*Strategy for handling model convergence issues in agonism analysis.* In several cases when modelling reproductive state-specific rates of agonistic interactions initiated or received, some candidate models (i.e., subsets of the ‘full’ model; see Methods) failed to converge, likely because of the model structure complexity relative to the infrequency of events. When this occurred, we followed a stepwise procedure. First, we rescaled numeric variables such that their mean values were centered at zero and their standard deviations were set to 1.000. If this failed to facilitate model convergence, we removed the fixed effects for rainfall and the quadratic group size term to simplify model structure. We chose rainfall because it was not a significant predictor of total agonistic interaction rate (model 1; $\beta = 3.1 \times 10^{-5}$, 95% CI = $[-4.9 \times 10^{-4}, 5.5 \times 10^{-4}]$). We chose the quadratic group size term because all significant quadratic relationships that we detected between group size and agonistic interaction rates reflected an initial decline in agonism with increasing group size and a subsequent flattening, rather than a true U- or inverted U-shaped relationship (Fig. S5). Finally, if some candidate models still failed to converge, we removed the random effect of group identity to simplify the random effects structure of the model. We chose group identity because it usually explained the least amount of total variance compared to individual identity or hydrological year.

Under sub-hypothesis 1, we constructed eight ‘full’ models predicting general agonistic interaction rates initiated toward and received from females in four reproductive states. All candidate models predicting the rate of agonistic interactions initiated toward females in early lactation converged without issue (Table S1; Supplementary Data; model 5). Rescaling numeric fixed effects resolved convergence issues for the models predicting the rate of agonistic interactions initiated toward and received from females in post-early lactation postpartum amenorrhea (models 9-10), received from cycling females (model 7), and initiated toward and received from pregnant females (models 6, 12). In order to resolve convergence issues for the models predicting the rate of agonistic interactions received from females in early lactation (model 8), we removed the rainfall and quadratic group size fixed effects, but retained the full random effects structure. To resolve convergence issues for the model predicting the rate of agonistic interactions initiated toward cycling females (model 11), we had to simplify the random effects structure by removing the random effect of group identity in addition to simplifying the fixed effects structure. Of the three models on which we base on our main claims (agonism initiated toward early lactating females, received from cycling females, and received from pregnant females; Fig. 2), none had a simplified fixed or random effects structure.

Under sub-hypothesis 2, we constructed four ‘full’ models predicting the rate at which early lactating females initiated agonistic interactions to females in four reproductive states, as a function of early lactational synchrony. Again, some candidate models failed to converge using our original model structure. Rescaling numeric fixed effects resolved convergence issues for the models predicting rates of agonistic interactions initiated by early lactating females to other early lactating females (model 14) and to females in post-early lactation postpartum amenorrhea (model 17). Rescaling numeric fixed effects and removing the fixed effects for rainfall and the group size quadratic term resolved convergence issues for the models predicting the rates that early lactating females initiated agonistic interactions toward cycling females (model 15) and pregnant females (model 16). To resolve convergence issues for the model predicting the rate that early lactating females initiated agonistic interactions toward all adult females (model 13), we also removed the random effect of group identity.

*Alternative hypothesis: generalized maternal aggression.* We hypothesized that the increase in total rates of agonistic interactions that females experience during early lactation was a result of heightened competition with other females for male social partners and food resources. However, this pattern could also reflect a broader mammalian trend for mothers to be more aggressive than other females as a direct strategy of infant defense[7-9]. To rule out this alternative hypothesis, we tested three of its predictions. First, if new mothers generally become more aggressive as an infant defense strategy, we predicted that females in early lactation would initiate agonistic interactions with other adult females more frequently than females in other reproductive states, regardless of the reproductive state of their targets. To test this prediction, we constructed a negative binomial GLMM with a log-link function predicting the number of times that an adult female initiated an agonistic interaction toward another adult female (of any reproductive state) within a focal sample (model 24, n = 60,655 focal samples). As fixed effects, we included (i) the focal female's proportional dominance rank, (ii) her reproductive state, (iii) her age, (iv) the number of nonfocal females present in the group, the total group size as a (v) linear and (vi) quadratic term, and (vii) the cumulative amount of rainfall that had fallen in the previous 30 days. We also included an offset term for the log-transformed number of minutes that the focal female was in sight of the observer during the focal sample and random effects for focal female identity, group identity, and hydrological year.

If female aggression primarily functions as infant defense, we also expected that females in early lactation would be more aggressive to the individuals most likely to cause their infants harm. In our analyses under sub-hypothesis 1, we found that the most frequent adult female aggressors of early lactating females (and, by extension, their clinging infants) were pregnant females. Thus, under this alternative hypothesis, we next predicted that females in early lactation would initiate agonistic interactions toward pregnant females more frequently than females in other reproductive states would. The analyses performed to test sub-hypothesis 1 in the main text provide a direct test of this prediction. They showed that reproductive state did not significantly predict the rate at which females initiated agonistic interactions toward pregnant females (model 12). Therefore, we did not pursue this prediction further.

Finally, because our results demonstrated that peripubertal females also pose a significant threat to the wellbeing of infants in the first 90 days of life, we also predicted that females in early lactation would initiate agonistic interactions toward peripubertal females at a rate higher than that of adult females in other reproductive states. To test this prediction, we again constructed a negative binomial GLMM with a log-link function, but with the number of times that an adult female initiated an agonistic interaction with a peripubertal female within a focal sample (model 25, n = 60,467 focal samples on 269 females where at least one nonfocal peripubertal female was present). We included the same fixed and random effects and offset term as in the model testing our first prediction under this hypothesis.

*Assigning causes of infant deaths.* For each infant that died, we attempted to categorize both the nature of the death (violence, pathology, or interruption of maternal care) and the agent of the death (i.e., what caused the violence, pathology, or interruption of maternal care) using all evidence available to us. We also assessed the confidence of both the nature and agent of the death, based on whether the cause was assigned from limited evidence that made the cause possible, circumstantial evidence that made the cause likely, or corroborating evidence that made the cause very likely or confirmed. Corroborating evidence for our highest level of confidence includes direct observations of death or events immediately preceding a presumed death, as well as observations of remains that are consistent with a particular death cause combined with further context that supports that cause. For example, in the case of infanticide, corroborating evidence may include direct observation of an infanticide, direct observation of an infant being attacked or kidnapped followed by the infant's disappearance, or observation of infant remains with infanticide-consistent wounds at particular times (e.g., after the immigration of swiftly rank-ascendant males, which sometimes perform infanticide in this population[10]).

*Impacts of ventral infant carrying on maternal foraging efficiency.* After finding support for a strong positive relationship between early lactational synchrony and the proportion of feeding time that low-ranking mothers spent in ventral contact with their infant (Table 3, Fig. 4e), we sought to investigate how a more restrictive maternal style might impact foraging efficiency in periods of intense female-female competition. Because baboons feed on many different food items that present different processing challenges[6], we anticipated that ventral infant presence would not impact all types of feeding to the same extent. Corm (the underground storage organ of some grass plants) feeding constitutes a major proportion of baboon feeding time in the long dry season, reaching an average peak of 0.594 ± 0.029 s.d. of total proportional feeding time in August[11]. Grass corms are characterized by a low energy yield rate relative to other foods in the baboon diet[6], but are an important fallback food when more preferred foods are not available. Extracting them from the dry ground requires cstrength and time, and female baboons often adopt a characteristic posture during corm feeding. Typically, they initially use one or both hands to partially expose the corm from the soil while seated. Next, they 'hunch' forward, crouched on their hind legs, to grab the corm with their teeth and/or hands before fully extracting it by rocking backward. In this posture, we expected that ventral infant presence would be a particular hindrance to efficient feeding.

To test this prediction, we first combined all baboon diet items into nine mutually exclusive categories, following[6,11,12]: grass corms, grass blades, grass blade bases and seedheads, *Vachellia* (formerly *Acacia*) gum, *Vachellia* blossoms, *Vachellia* seeds, fruits, animal matter, and 'other'. Because focal females can switch food items within a single focal sample, we focused on the first minute of each focal sample and identified the focal female's instantaneous activity, posture (seated vs. standing), and infant contact state (ventral, dorsal, or neither), and near neighbor status (adult male neighbor present vs. absent).

We used these data to construct four Bernoulli GLMMs with logit-link functions. Using all data when adult females were feeding on the first minute of their focal sample (model 28; n = 23,170 behavioral records on 268 females), we first constructed a model predicting whether or not the feeding focal female was standing. As fixed effects, we included: (i) the category of food she was eating, (ii) her 'minor' reproductive state, (iii-iv) the size of her group as a linear and quadratic term, (v) her ordinal dominance rank, (vi) her age, and (vii) the cumulative amount of rainfall that had fallen in the previous 30 days. We also included focal female identity, group identity, and hydrological year as random effects.

Our next model used only data on feeding females in early lactation (model 29; n = 2,709 feeding records on 231 females) and the response variable was whether or not the focal female was in ventral contact with her infant. Food category was the primary fixed effect of interest, but we included additional controls for (ii) male neighbor presence, (iii) focal female ordinal dominance rank, (iv) focal female age, (v) group size, and (vi) recent cumulative rainfall, as in model 2. Again, we included female identity, group identity, and hydrological year as random effects.

After finding strong evidence that corm feeding was largely incompatible with ventral infant presence (see Supplementary Results), we tested whether low-ranking females experienced a synchrony-related energetic cost in the form of reduced corm feeding. We used all behavioral data from the first minute of focal samples on early lactating females (model 30; n = 7,373 behavioral records on 235 females) to construct a GLMM predicting whether or not the focal female was feeding on grass corms as a function of the interaction between her dominance rank and early lactational synchrony, analogous to model 24 (Fig. 4e). The fixed effects were (i) early lactational synchrony, (ii) the focal female's ordinal dominance rank, (iii) the interaction between (i) and (ii), (iv) the focal female's age, the focal female's group size as a (v) linear and (vi) quadratic term, and (vii) recent cumulative rainfall. Female identity, group identity, and hydrological year were included as random effects.

Finally, we thought low-ranking females might compensate for potential reductions in corm feeding during periods of high early lactational synchrony by increasing feeding time spent on non-corm foods. Therefore, we constructed a GLMM predicting whether or not a focal early lactating female was was feeding on a non-corm food on the first minute of each focal sample (model 31; n = 7,373 behavioral records on 235 females). We included the same fixed and random effects as in model 30.

*Peripubertal female neighbors.* We performed a *post hoc* analysis to assess if infanticides were facilitated by increased infant exposure to peripubertal females during times of high early lactational synchrony. Specifically, using focal follow data collected on adult females in early lactation, we constructed two Bernoulli GLMMs with a logit link function predicting the probability that the nearest neighbor within 5 m of an early lactating mother was a peripubertal female. We considered peripubertal females to be those who (i) were at least 3 years old (the youngest age at which a female was observed to commit a fatal kidnapping), (ii) were younger than 8 years old (the oldest approximate age at which a female might be expected to have her first live birth), and (iii) had never given birth to a live offspring. Because peripubertal females may have greater agency to kidnap the infant of a relatively lower-ranking female than the infant of a higher-ranking female, we separately modelled (i) the probability that the nearest neighbor of an early lactating mother was a higher-ranking peripubertal female (relative to the mother; model 26) and (ii) the probability that the nearest neighbor of an early lactating mother was a lower-ranking peripubertal female (again, relative to the mother's own rank; model 27). As fixed effects in both models, we included (i) the total number of peripubertal females present in the group who were higher- or lower-ranking than the focal female, which controls for their opportunity to appear as a nearest neighbor and (ii) the proportion of adult females in the group who were in the first 90 days of lactation. We also included focal female identity, group identity, and hydrological year as random effects to control for repeated sampling of mothers and groups.

In our binomial GLMMs predicting association time with adult males, we modelled the proportion of observation time spent within 5 m of at least one adult male as our response variable. Due to the nature of our focal sampling protocol (see above), we can determine whether or not an adult male was in close proximity to a focal female at every minute of a given sample (and thus calculate a total proportion of association time for each sample). However, because peripubertal females are only recorded as neighbors when they are the *nearest* neighbor of a focal female, it is impossible to determine whether or not a peripubertal female was in close proximity to a focal female at every minute of a given sample. Therefore, in these *post hoc* neighbor analyses, we instead modelled the probability that a focal female's nearest neighbor was a peripubertal female in the first minute of each focal sample. We also excluded any focal samples collected when either no higher- or lower-ranking peripubertal females were present in the group. Our final dataset for the higher-ranking neighbor model included 5,846 observations on 210 early lactating females in 14 social groups from October 1999 to December 2021 and our final dataset for the lower-ranking neighbor model included 7,059 observations on 228 females in 14 social groups across the same time window.

**Supplementary Results & Discussion**

*Validating the 90-day threshold of early lactation*. When we visualized changes in the four competition-related behaviors of interest as functions of days since parturition, it was apparent that mothers underwent detectable decreases or increases in each behavioral metric over the first year of postpartum amenorrhea (Fig. S1-2). A major shift in the mean proportion of time that females spent in close proximity to adult males corresponded almost exactly with our expected 90-day threshold (Fig. S1c, S2c). Specifically, in the first 90 days after giving birth, mothers maintained proximity with males for a consistently high proportion of their active hours. At 90 days postpartum, the time that females spent in association with males started to decline, and this decline was sustained throughout the remainder of their infant's first year of life. Similarly, maternal feeding time spent in ventral contact with an infant was higher in the first 90 days postpartum and lower after this period, but this decline started slightly before the first decline in time spent with males (Fig. S1a, S2a). Agonistic interaction rates were also higher in the first 90 days postpartum than after (Fig. S2d), but these data were considerably noisier than the consistent shifts in time with males and feeding time with a ventral infant (Fig. S1d). Females spent very little time foraging immediately after giving birth (Fig. S1b, Fig. S2b). Foraging time showed a steep increase from the first 10-day interval postpartum to the second, after which it showed a steady increase over the next year.

*Alternative hypothesis: maternal aggression.* We did not find support for the alternative hypothesis that increased total rates of agonistic interaction during early lactation were the result of maternal aggression for the purpose of infant defense. Females in early lactation did not generally initiate agonistic interactions with other adult females more frequently than females in other reproductive states (model 24). Females in early lactation also did not initiate more agonistic interactions with the specific classes of females who posed the highest risks to their infants (pregnant females, model 12, or peripubertal females, model 25). Rather, when females in early lactation initiated agonstic interactions, they tended to target other females in early lactation (model 5, Fig. 2a), in support of the hypothesis that aggression by early lactating females is a response to competition over male social partners.

*Relationship between ventral infant carrying and maternal foraging efficiency*. In support of the idea that mothers of the youngest infants have reduced mobility while foraging, early lactating females were significantly less likely to be standing while feeding than females in any other minor reproductive state (Supplementary Data, model 28). For example, when feeding on grass blades, the predicted probability of standing was 0.311 (95% CI: [0.261, 0.367]) for early lactating females, but this value ranged from 0.351-0.610 in females in other reproductive states. Food category also predicted the probability of standing while feeding. As expected, the probability of standing during grass corm feeding was significantly lower than when feeding on any other food category (model 28). When feeding on grass corms, early lactating females had a predicted probability of standing of 0.080 (95% CI: [0.064, 0.099], compared to 0.311-0.473 for other food categories. This difference reflects the fact that corm extraction imposes greater postural constraints on female baboons than other types of feeding do.

Early lactating female behavior suggested that ventral infant presence presents a challenge for corm feeding, but not for feeding on other foods. When feeding on corms, early lactating females were significantly less likely to be in ventral contact with their infant than when feeding on any other food category (model 29; Fig. S4a). New mothers had a predicted probability of ventral infant contact during corm feeding of 0.063 (95% CI: [0.048, 0.082]), and a predicted range of 0.190-0.370 across other food types. When these mothers fed on above-ground grass parts (blades, blade bases, and seedheads), they were 3.2x to 4.1x more likely to be in ventral contact with their infant than when they fed on grass corms. Ventral infant presence seemed particularly incompatible with seated corm feeding (which represents the vast majority of corm feeding records). Across 1,130 seated corm feeding records, early lactating females were in ventral contact with their infant 0.708% of the time, compared to 59.633% of 110 standing corm feeding records. This suggests that ventral infant presence interferes with efficient corm feeding.

As early lactational synchrony increases, low-ranking mothers increased the proportion of feeding time they spent in ventral contact with their infant (Fig. 4e), potentially as a protective measure in response to reduced access to male protective services (Fig. 4a). Given that clinging infants likely interfere with corm feeding, low-ranking females in periods of high synchrony may not be able to easily access this critical fallback food without leaving their infant vulnerable to negative conspecific attention. Models 30-31 imply that this tradeoff (between foraging efficiency and infant safety) imposes an energetic cost on low-ranking mothers. First, while high-ranking females maintained or slightly increased their time spent feeding on corms as early lactational synchrony increased, low-ranking females experienced a strong decline in corm feeding time with increasing early lactational synchrony (model 30, ordinal dominance rank: $\beta = 0.040$, 95% CI: [0.016, 0.065]; early lactational synchrony: $\beta = 0.877$, 95% CI: [-0.189, 1.943]; interaction term: $\beta = -0.188$, 95% CI: [-0.294, -0.083]; Fig. S4b). On a day with no recent rainfall (when corm feeding is most prevalent) at the median level of early lactational synchrony (0.105), the lowest-ranking early lactating females were predicted to spend 32.847% of their total time feeding on grass corms. In comparison, on a day with no recent rainfall at our high synchrony benchmark (0.320), the lowest-ranking early lactating females were predicted to spend 15.433% of their total time feeding on grass corms. This suggests that, on the most synchronous days, the lowest-ranking mothers intake approximately half the amount of grass corms that they would on an average day.

Low-ranking mothers may compensate for this shortfall by increasing consumption of non-corm foods when early lactational synchrony is high. However, model 31 suggests otherwise. The proportion of time that early lactating females spent feeding on non-corm foods was not predicted by rank, early lactational synchrony, or the interaction between rank and early lactational synchrony (model 31, ordinal dominance rank: $\beta$ = -0.035, 95% CI: [-0.082, 0.013]; early lactational synchrony: $\beta$ = 0.366, 95% CI: [-1.397, 2.128]; interaction term: $\beta$ = 0.132, 95% CI: [-0.051, 0.314; Fig. S4c]). Together, these results imply that early lactational synchrony may be particularly energetically costly for low-ranking mothers when synchrony occurs in the long dry season because the increased need to protect vulnerable infants limits mothers' access to the primary component of the dry season diet.

*Peripubertal female neighbors*. As early lactational synchrony increased, an early lactating female was less likely to have a higher-ranking peripubertal female as her nearest neighbor (model 32, GLMM: $\beta$ = -1.324, 95% CI: [-2.409, -0.240]). Early lactational synchrony had no association with the probability that an early lactating female's nearest neighbor was a lower-ranking peripubertal female (model 33, GLMM: $\beta$ = -0.141, 95% CI: [-0.822, 0.540]).

### Supplementary Tables

*Table S1. Summary of all 'full' statistical models*. Each row describes the model structure of a 'full' statistical model (i.e., including all hypothesized covariates; see Methods for detailed rationale on covariate inclusion and model selection) described in the manuscript main text. Models 1-24 and 26-36 were generalized linear mixed models and models 25 and 37-39 were Cox proportional hazards models. Fixed effects that were found to be significantly correlated with the response variable are indicated in bold. See table footnotes for explanations of abbreviations used in the 'Fixed effects' and 'Random effects' columns.

| Sub-hypothesis | Model | Focal subjects | Response variable | Fixed effects | Random effects |
|---|---|---|---|---|---|
| 1. Baboons in early lactation are focal points for female-female competition | 1 | adult females | total agonistic interaction rate | **ReproStateMin** + **Rank** + **Rank$^2$** + **Age** + **GrpSize** + **GrpSize$^2$** + Rainfall + **CountFemale** + Offset | Female ID + Group ID + HydroYear |
| | 2 | adult females | proportion of feeding time spent in ventral contact with an infant | **ReproStateMin** + Rank + Age + **GrpSize** + **Rainfall** + **PropTimeMale** | Female ID + Group ID + HydroYear |
| | 3 | adult females | proportion of time spent foraging | **ReproStateMin** + Rank + **Age** + **GrpSize** + **GrpSize$^2$** + **Rainfall** | Female ID + Group ID + HydroYear |
| | 4 | adult females | proportion of time spent close to an adult male | **ReproStateMin** + **Rank** + **Age** + **Rainfall** + **CountMale** + **CountCycling** | Female ID + Group ID + HydroYear |
| | 5 | adult females | rate of agonistic interactions initiated toward early lactating females | **ReproStateMaj** + **Rank** + **Age** + **GrpSize** + **GrpSize$^2$** + Rainfall + **CountEarlyLact** + Offset | Female ID + Group ID + HydroYear |
| | 6 | adult females | rate of agonistic interactions received from pregnant females | **ReproStateMaj** + **Rank** + Age + **GrpSize** + **GrpSize$^2$** + Rainfall + **CountPregnant** + Offset | Female ID + Group ID + HydroYear |
| | 7 | adult females | rate of agonistic interactions received from cycling females | **ReproStateMaj** + **Rank** + **Age** + **GrpSize** + GrpSize$^2$ + Rainfall + **CountCycling** + Offset | Female ID + Group ID + HydroYear |
| | 8* | adult females | rate of agonistic interactions received from early lactating females | ReproStateMaj + **Rank** + Age + **GrpSize** + **CountEarlyLact** + Offset | Female ID + Group ID + HydroYear |
| | 9 | adult females | rate of agonistic interactions initiated | ReproStateMaj + **Rank** + **Age** + GrpSize + GrpSize$^2$ + Rainfall + **CountPostEL** + Offset | Female ID + Group ID + HydroYear |

| | | | | | |
|---|---|---|---|---|---|
| | | | toward post-early lactation postpartum amenorrhea females | | |
| | 10 | adult females | rate of agonistic interactions received from post-early lactation postpartum amenorrhea females | ReproStateMaj + **Rank** + Age + **GrpSize** + GrpSize$^2$ + Rainfall + **CountPostEL** + Offset | Female ID + Group ID + HydroYear |
| | 11* | adult females | rate of agonistic interactions initiated toward cycling females | ReproStateMaj + **Rank** + **Age** + **GrpSize** + **CountCycling** + Offset | Female ID + HydroYear |
| | 12 | adult females | rate of agonistic interactions initiated toward pregnant females | ReproStateMaj + **Rank** + Age + **GrpSize** + GrpSize$^2$ + Rainfall + **CountPregnant** + Offset | Female ID + Group ID + HydroYear |
| 2. Female-female competition intensifies when early lactational synchrony is high | 13* | early lactating females | rate of agonistic interactions intiated toward adult females | **EarlyLactSynch** + **Rank** + Age + **GrpSize** + **CountFemales** + Offset | Female ID + HydroYear |
| | 14 | early lactating females | rate of agonistic interactions intiated toward early lactating females | EarlyLactSynch + **Rank** + Age + GrpSize + GrpSize$^2$ + Rainfall + CountEarlyLact + Offset | Female ID + Group ID + HydroYear |
| | 15* | early lactating females | rate of agonistic interactions intiated toward cycling females | **EarlyLactSynch** + **Rank** + **Age** + **GrpSize** + **CountCycling** + Offset | Female ID + Group ID + HydroYear |
| | 16* | early lactating females | rate of agonistic interactions intiated toward pregnant females | EarlyLactSynch + **Rank** + **Age** + GrpSize + **CountPregnant** + Offset | Female ID + Group ID + HydroYear |
| | 17 | early lactating females | rate of agonistic interactions intiated toward post-early lactation postpartum amenorrhea females | EarlyLactSynch + **Rank** + Age + GrpSize + GrpSize$^2$ + **Rainfall** + CountPostEL + Offset | Female ID + Group ID + HydroYear |
| | 18 | early lactating females | proportion of time spent close to an adult male | EarlyLactSynch + **Rank** + (**EarlyLactSynch x Rank)** + Age + Rainfall + **CountMale** + **CountCycling** | Female ID + Group ID + HydroYear |

| | | | | | |
|---|---|---|---|---|---|
| | 19 | pregnant females | proportion of time spent close to an adult male | **EarlyLactSynch** + **Rank** + (**EarlyLactSynch x Rank)** + Age + **Rainfall** + **CountMale** + **CountCycling** | Female ID + Group ID + HydroYear |
| | 20 | early lactating females | proportion of time spent close to an adult male | EarlyLactSynch + Rank + **(EarlyLactSynch x Rank)** + Age + Rainfall + **CountMale** + **(EarlyLactSynch x CountMale)** + **CountCycling** | Female ID + Group ID + HydroYear |
| | 21 | pregnant females | proportion of time spent close to an adult male | **EarlyLactSynch** + **Rank** + **(EarlyLactSynch x Rank)** + Age + **Rainfall** + **CountMale** + (EarlyLactSynch x CountMale) + **CountCycling** | Female ID + Group ID + HydroYear |
| | 22 | early lactating females | proportion of time spent foraging | EarlyLactSynch + Rank + **(EarlyLactSynch x Rank)** + Age + **Rainfall** + GrpSize + $\text{GrpSize}^2$ | Female ID + Group ID + HydroYear |
| | 23 | pregnant females | proportion of time spent foraging | EarlyLactSynch + Rank + **(EarlyLactSynch x Rank)** + Age + **Rainfall** + **GrpSize** + **$\text{GrpSize}^2$** | Female ID + Group ID + HydroYear |
| | 24 | early lactating females | proportion of feeding time spent in ventral contact with an infant | **EarlyLactSynch** + **Rank** + **(EarlyLactSynch x Rank)** + Age + **Rainfall** + GrpSize | Female ID + Group ID + HydroYear |
| 3. Female-female competition related to high early lactational synchrony increases infant mortality risk | 25 | infants | mortality risk | **EarlyLactSynch** + **MaternalDeath** + **MaternalRank** + MaternalAge + $\text{MaternalAge}^2$ + GrpSize + Rainfall + tt(MaternalDeath) + tt(MaternalAge) + tt($\text{MaternalAge}^2$) + tt(GrpSize) | Infant ID (cluster term) |
| HA: Generalized maternal aggression | 26 | adult females | rate of agonistic interactions initiated toward adult females | ReproStateMaj + **Rank** + **Age** + **GrpSize** + **$\text{GrpSize}^2$** + Rainfall + **CountFemale** + Offset | Female ID + Group ID + HydroYear |
| | 27 | adult females | rate of agonistic interactions initiated toward peripubertal females | ReproStateMaj + **Rank** + **Age** + GrpSize + $\text{GrpSize}^2$ + Rainfall + **CountPeri** + Offset | Female ID + Group ID + HydroYear |
| *Post-hoc* analyses: | 28 | adult females | proportion of feeding records spent standing | **FoodCat** + **ReproStateMin** + **GrpSize** + **$\text{GrpSize}^2$** + Rank + **Age** + **Rainfall** | Female ID + Group ID + HydroYear |

| | | | | | |
|---|---|---|---|---|---|
| impacts of ventral infant carrying on maternal foraging efficiency | 29 | early lactating females | proportion of feeding records spent in ventral contact with an infant | **FoodCat** + Rainfall + MaleNeigh + GrpSize + Rank + Age | Female ID + Group ID + HydroYear |
| | 30 | early lactating females | proportion of time spent feeding on grass corms | EarlyLactSynch + **Rank + (EarlyLactSynch x Rank)** + **Rainfall** + Age + GrpSize + $GrpSize^2$ | Female ID + Group ID + HydroYear |
| | 31 | early lactating females | proportion of time spent feeding on non-grass corm foods | EarlyLactSynch + Rank + (EarlyLactSynch x Rank) + **Rainfall** + Age + GrpSize + $GrpSize^2$ | Female ID + Group ID + HydroYear |
| *Post-hoc* analyses: proximity to peripubertal females (potential kidnappers) | 32 | early lactating females | probability that nearest neighbor within 5 m is a higher-ranking peripubertal female | **EarlyLactSynch** + **CountHigherPeri** | Female ID + Group ID + HydroYear |
| | 33 | early lactating females | probability that nearest neighbor within 5 m is a lower-ranking peripubertal female | EarlyLactSynch + **CountLowerPeri** | Female ID + Group ID + HydroYear |
| Alternate metrics of lactational synchrony | 34* | early lactating females | rate of agonistic interactions intiated toward adult females | **EarlyMidLactSynch** + **Rank** + Age + GrpSize + CountFemale + Offset | Female ID + HydroYear |
| | 35* | early lactating females | rate of agonistic interactions intiated toward adult females | LateMidLactSynch + **Rank** + Age + GrpSize + CountFemale + Offset | Female ID + HydroYear |
| | 36* | early lactating females | rate of agonistic interactions intiated toward adult females | LateLactSynch + **Rank** + Age + GrpSize + CountFemale + Offset | Female ID + HydroYear |
| | 37 | infants | mortality risk | EarlyMidLactSynch + **MaternalDeath** + **MaternalRank** + MaternalAge + $MaternalAge^2$ + **GrpSize** + Rainfall + tt(MaternalDeath) + tt(MaternalAge) + tt($MaternalAge^2$) + tt(GrpSize) | Infant ID (cluster term) |
| | 38 | infants | mortality risk | LateMidLactSynch + **MaternalDeath** + **MaternalRank** + MaternalAge + $MaternalAge^2$ + **GrpSize** + Rainfall + tt(MaternalDeath) + tt(MaternalAge) + tt($MaternalAge^2$) + tt(GrpSize) | Infant ID (cluster term) |
| | 39 | infants | mortality risk | LateLactSynch + **MaternalDeath** + **MaternalRank** + MaternalAge + $MaternalAge^2$ + **GrpSize** + | Infant ID (cluster term) |

| | |
|---|---|
| | Rainfall + tt(MaternalDeath) + tt(MaternalAge) + tt($MaternalAge^2$) + tt(GrpSize) |

*Model structure was simplified to facilitate convergence (see Supplementary Methods for more information)

CountCycling = number of nonfocal cycling females present

CountEarlyLact = number of nonfocal early lactating females present

CountFemale = number of nonfocal adult females present

CountHigherPeri = number of nonfocal, higher-ranking peripubertal females present

CountLowerPeri = number of nonfocal, lower-ranking peripubertal females present

CountMale = number of adult males present

CountPeri = number of nonfocal peripubertal females present

CountPostEL = number of nonfocal post-early lactation postpartum amenorrhea females present

CountPregnant = number of nonfocal pregnant females present

EarlyLactSynch = early lactational synchrony

EarlyMidLactSynch = early-mid lactational synchrony

FoodCat = category of food that the focal female was currently consuming (reference category = grass corms)

GrpSize = total group size

HydroYear = hydrological year

LateLactSynch = late lactational synchrony

LateMidLactSynch = late-mid lactational synchrony

MaleNeigh = whether or not the focal female was currently within 5 m of an adult male

MaternalAge = mother’s age on the day of the infant’s birth

MaternalDeath = whether mother was dead or alive on a given day

MaternalRank = mother’s dominance rank on the day of the infant’s birth

Offset = offset term term for the log-transformed number of minutes that the focal female was in sight of the observer during the focal sample

PropTimeMale = proportion of minutes in the focal sample at which the focal female was within 5 m of at least one adult male

Rank = dominance rank

ReproStateMaj = ‘major’ reproductive state (reference category = early lactation)

ReproStateMin = ‘minor’ reproductive state (reference category = early lactation)
tt() = time transformation term

*Table S2. Group-level information on the mean frequency and duration of periods of 'high' early lactational synchrony (ELS)*. A period of 'high' early lactational synchrony within a group was defined as a period of time when the proportion of adult females who were in early lactation remained at or above 0.320 (the 95th percentile value overall).

| Group ID | Observation time (years) | Number of ELS bouts | Mean ELS bout duration (days) | Mean frequency of ELS bouts (year $^{-1}$) |
|---|---|---|---|---|
| **1** | 16.427 | 13 | 30.846 | 0.791 |
| **2** | 17.095 | 13 | 22.846 | 0.760 |
| **3** | 11.146 | 6 | 15.000 | 0.538 |
| **4** | 14.856 | 10 | 22.000 | 0.673 |
| **5** | 14.869 | 14 | 24.857 | 0.942 |
| **6** | 1.692 | 0 | – | 0.000 |
| **7** | 3.806 | 2 | 10.500 | 0.526 |
| **8** | 9.481 | 5 | 25.200 | 0.527 |
| **9** | 5.873 | 3 | 6.667 | 0.511 |
| **10** | 4.145 | 2 | 45.500 | 0.482 |
| **11** | 12.230 | 12 | 26.667 | 0.981 |
| **12** | 10.642 | 1 | 16.000 | 0.094 |
| **13** | 4.093 | 5 | 37.800 | 1.222 |
| **14** | 2.138 | 2 | 59.000 | 0.935 |
| **15** | 13.769 | 9 | 8.444 | 0.654 |
| **16** | 3.184 | 2 | 71.000 | 0.628 |
| **17** | 0.523 | 1 | 4.000 | 1.912 |
| **18** | 13.133 | 6 | 50.667 | 0.457 |
| **19** | 0.575 | 0 | – | 0.000 |

*Table S3. All cases of observed or possible infanticide of infants less than 90 days old in the study population between 1976 and 2021* (the period covered by our survival analysis). The eight infant deaths that occurred on days of 'high' early lactational synchrony (i.e., days exceeding the mean annual maximum value of 32.0% of females in early lactation, which corresponds to the 95th percentile value overall) are highlighted in bold; of these, six were known or strongly suspected to have been caused by females. Infanticide events included in the Fisher's exact test (i.e., those that were directly observed or supported by corroborating evidence) are marked with an asterisk.

| Case number | Season-Year | Proportion of adult females in early lactation | Group size | Actor sex | Type | Context | Evidence |
|---|---|---|---|---|---|---|---|
| **1*** | **Wet 1979** | **0.643** | **46** | **Female** | **Kidnapping, no severe aggression** | **Intragroup** | **Direct observation of kidnapping; described in[13]** |
| **2** | Wet 1980 | Unknown | 32 | Mixed sex group | Severe aggression | Intergroup | Direct observation of kidnapping and severe multiparty attack; described in[14] |
| **3** | Wet 1981 | 0.231 | 32 | Male (likely) | Unknown | Intragroup (likely) | Circumstantial; remains observed; recent arrival of rank-rising male[10] |
| **4** | Wet 1990 | 0.238 | 64 | Unknown | Severe aggression | Unknown | Circumstantial; remains observed |
| **5** | Dry 1995 | 0.143 | 24 | Unknown | Severe aggression | Unknown | Circumstantial; remains observed |
| **6** | Wet 1997 | 0.222 | 26 | Female | Kidnapping, no severe aggression | Intragroup | Direct observation of kidnapping |
| **7*** | **Wet 2004** | **0.357** | **43** | **Female** | **Kidnapping, no severe aggression** | **Intragroup** | **Direct observation of kidnapping** |
| **8** | **Wet 2004** | **0.381** | **71** | **Unknown, possibly female** | **Unknown** | **Unknown, possibly intragroup** | **Limited; remains observed; preceded by high frequency of multiparty female-female aggression** |
| **9** | Wet 2005 | 0.286 | 46 | Female | Kidnapping, no severe aggression | Intragroup | Direct observation of kidnapping |

| | | | | | | | |
|---|---|---|---|---|---|---|---|
| **10** | Wet 2006 | 0.267 | 48 | Female | Kidnapping, no severe aggression | Intragroup | Direct observation of kidnapping |
| **11*** | **Dry 2010** | **0.500** | **101** | **Female (likely)** | **Severe aggression (likely)** | **Intragroup (likely)** | **Corroborating; mother observed without infant and with severe injuries; occurred during an intense period of severe female aggression targeted at high-ranking females and their offspring** |
| **12*** | **Dry 2010** | **0.500** | **100** | **Female** | **Kidnapping, no severe aggression** | **Intragroup** | **Direct observation of actor holding corpse; occurred during an intense period of severe female aggression targeted at high-ranking females and their offspring** |
| **13*** | **Dry 2010** | **0.500** | **99** | **Female (likely)** | **Severe aggression (likely)** | **Intragroup (likely)** | **Corroborating; infant was first noted missing when mother was the target of severe multiparty female aggression; occurred during an intense period of severe female aggression targeted at high-ranking females and their offspring** |
| **14** | Wet 2011 | 0.0769 | 45 | Unknown | Unknown | Unknown | Limited; remains observed |
| **15** | Dry 2012 | 0.25 | 34 | Male | Severe aggression; cannibalism | Intragroup | Direct observation of cannibalism by actor |
| **16** | Wet 2013 | 0.214 | 51 | Male | Kidnapping, no severe aggression | Intragroup | Direct observation of kidnapping |
| **17** | Wet 2013 | 0.214 | 50 | Male | Kidnapping, no severe aggression | Intragroup | Direct observation of kidnapping |
| **18** | Wet 2013 | 0.214 | 50 | Male (likely) | Kidnapping, no severe aggression | Intragroup (likely) | Corroborating; remains observed; recent arrival of rank-rising male[10] |
| **19** | Wet 2014 | 0.0588 | 53 | Male | Kidnapping, no severe aggression | Intragroup | Direct observation of kidnapping |
| **20** | Dry 2015 | 0.25 | 40 | Female | Kidnapping, no severe aggression | Intragroup | Direct observation of kidnapping |
| **21** | Wet 2016 | 0.0588 | 50 | Unknown | Kidnapping, no severe aggression (likely) | Intragroup (likely) | Limited; remains observed |

| | | | | | | | |
|---|---|---|---|---|---|---|---|
| **22** | Wet 2017 | 0.0588 | 49 | Female | Kidnapping, no severe aggression | Intragroup | Direct observation of kidnapping |
| **23** | **Wet 2018** | **0.500** | **21** | **Unknown** | **Kidnapping, no severe aggression (likely)** | **Intragroup (likely)** | **Limited; remains observed** |
| **24** | Wet 2019 | 0.292 | 78 | Unknown | Unknown | Unknown | Limited; no remains observed |
| **25** | Wet 2020 | 0.167 | 50 | Female | Kidnapping, no severe aggression | Intragroup | Direct observation of actor holding corpse |
| **26** | Dry 2020 | 0.227 | 61 | Female | Kidnapping, no severe aggression | Intragroup | Direct observation of kidnapping |
| **27*** | **Dry 2020** | **0.471** | **57** | **Female** | **Kidnapping, no severe aggression** | **Intragroup** | **Direct observation of kidnapping** |
| **28** | Wet 2021 | 0.136 | 70 | Female | Kidnapping, no severe aggression | Intergroup | Direct observation of kidnapping |

*Table S4. Sample sizes for behavioral data on females in each reproductive state.*

| 'Major' reproductive state | 'Minor' reproductive state | Number of focal samples | Number of observation minutes |
|---|---|---|---|
| Postpartum amenorrhea | Early lactation | 7,640 | 73,186 |
| | Early-mid lactation | 6,674 | 63,823 |
| | Late-mid lactation | 5,470 | 52,266 |
| | Late lactation | 3,230 | 30,745 |
| | Post-year postpartum amenorrhea | 1,657 | 15,816 |
| | **Total** | **24,671** | **235,836** |
| Ovarian cycling | Flat | 3,870 | 36,973 |
| | Turgescence-follicular | 7,494 | 71,598 |
| | Turgescence-ovulation | 2,587 | 24,722 |
| | Deturgescence-luteal | 5,052 | 48,291 |
| | **Total** | **19,003** | **181,584** |
| Pregnancy | First trimester | 5,939 | 57,030 |
| | Second trimester | 5,736 | 54,983 |
| | Third trimester | 5,316 | 51,093 |
| | **Total** | **16,991** | **163,106** |
| All adult females | **Total** | **60,665** | **580,526** |

*Table S5. Results of the 'full' model predicting infant mortality risk in the first 90 days of life* (Cox proportional hazards model; total infants = 1,304; died = 140, censored = 33). Covariates where the 95% CI did not overlap zero are indicated in bold. Model predictions are presented to aid interpretation of estimates and were calculated for the extreme values of each significant fixed effect, while holding values for other fixed effects constant at their observed mean value.

| Fixed effects | Estimate | Std. Error | 95 % CI | Hazard ratio | Model predictions |
|---|---|---|---|---|---|
| **Maternal death** | **7.026** | **2.190** | **[1.705, 12.347]** | **1125.816** | When an infant's mother is dead, mortality risk is 1125.452 times higher than when she is alive |
| tt(Maternal death) | $-3.2 \times 10^{-4}$ | $9.2 \times 10^{-4}$ | [-0.003, 0.002] | 1.000 | – |
| **Early lactational synchrony** | **2.975** | **0.767** | **[1.466, 4.484]** | **19.590** | At max. ELS (0.712), mortality risk is 7.607 times higher than at min. ELS (0.032) |
| Maternal age | -0.230 | 0.123 | [-0.469, 0.009] | 0.794 | – |
| tt(Maternal age) | -0.001 | 0.002 | [-0.004, 0.001] | 0.999 | – |
| **Maternal age$^2$** | **0.010** | **0.005** | **[$2.6 \times 10^{-4}$, 1.930]** | **1.010** | At younget maternal age (4.8 y), mortality risk is 1.587 times higher than at median maternal age (10.1 y)<br><br>At oldest maternal age (23.9 y), mortality risk is 4.064 times higher than at median maternal age (10.1 y) |
| tt(Maternal age$^2$) | – | – | – | – | – |
| **Maternal rank** | **0.036** | **0.016** | **[0.005, 0.067]** | **1.036** | At lowest maternal rank (29), mortality risk is 2.716 times higher than at highest maternal rank (1) |
| Rainfall | $-3.8 \times 10^{-4}$ | 0.002 | [-0.004, 0.003] | 1.000 | – |
| Group size | -0.009 | 0.006 | [-0.020, 0.002] | 0.991 | – |
| tt(Group size) | -0.005 | 0.009 | [-0.019, 0.009] | 0.995 | – |

*Table S6. Sample sizes of infant death causes.*

| Cause | All infant deaths | Infant deaths above the 95$^{th}$ percentile value of ELS | Infant deaths below the 95$^{th}$ percentile value of ELS |
|---|---|---|---|
| Unknown | 95 | 18 | 77 |
| Infanticide by a female | 14 | 6 | 8 |
| Infanticide by a male | 5 | 0 | 5 |
| Noninfectious pathology | 18 | 2 | 16 |
| Infectious pathology | 2 | 0 | 2 |
| Maternal death | 2 | 0 | 2 |
| Interruption of maternal care | 3 | 1 | 2 |
| Human/domestic dog | 1 | 0 | 1 |

**Supplementary Figures**

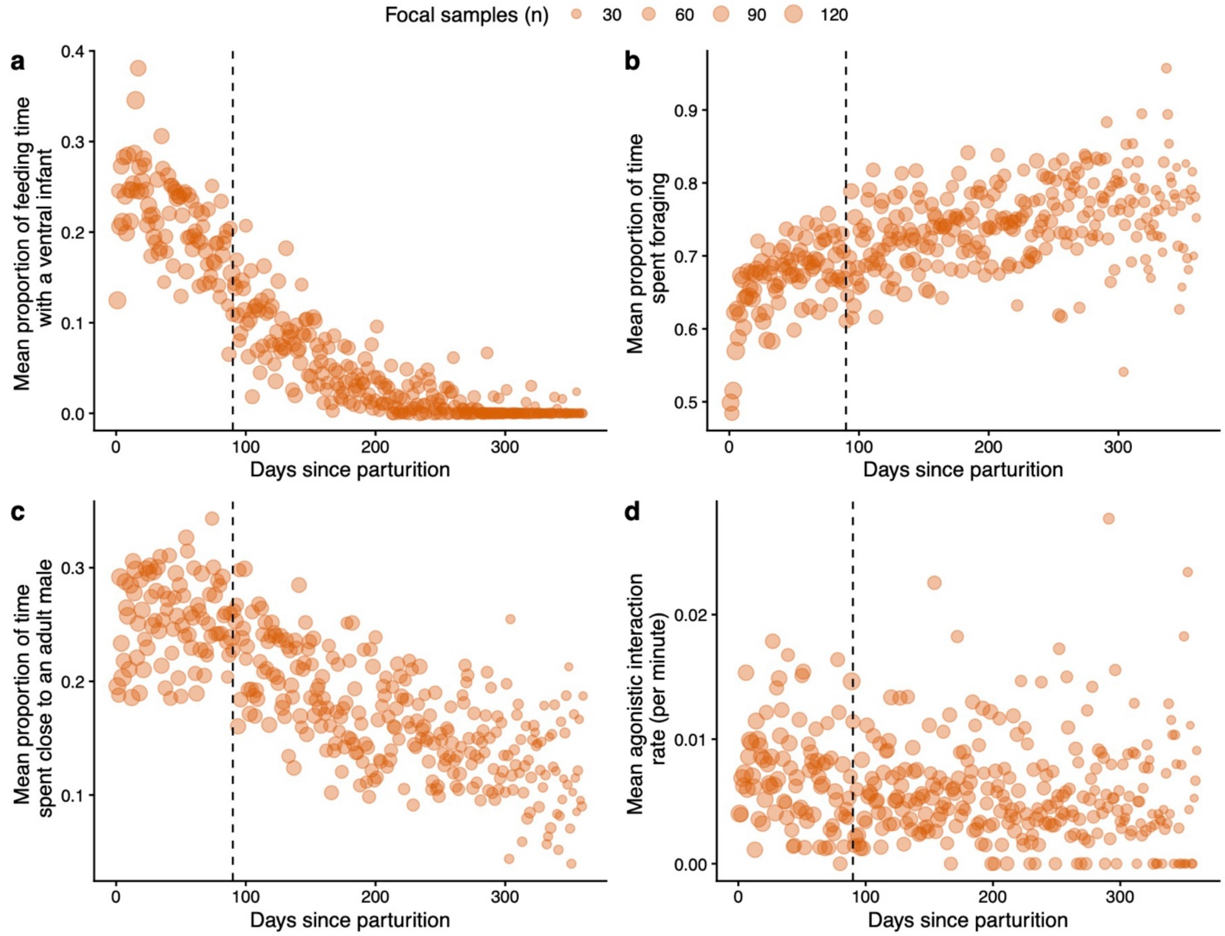

*Figure S1. The first 90 days postpartum are characterized by a dual challenge to mothers that affects both time budgets and behavior (daily values).* In this time period, mothers spent (a) a greater proportion of feeding time in ventral contact with an infant, (b) a much lower proportion of time foraging, and (c) a greater proportion of time within 5 m of adult males than in later postpartum periods. (d) They also had higher rates of female-female agonistic interactions in the first 90 days postpartum than in later postpartum periods. Each orange circle represents the mean value across all focal samples conducted on a given day since parturition, scaled proportionally in size to the number of focal samples in our data set which were conducted on that day. The dotted vertical line is placed at 90 days postpartum, our defined threshold demarcating early lactation from the rest of postpartum amenorrhea.

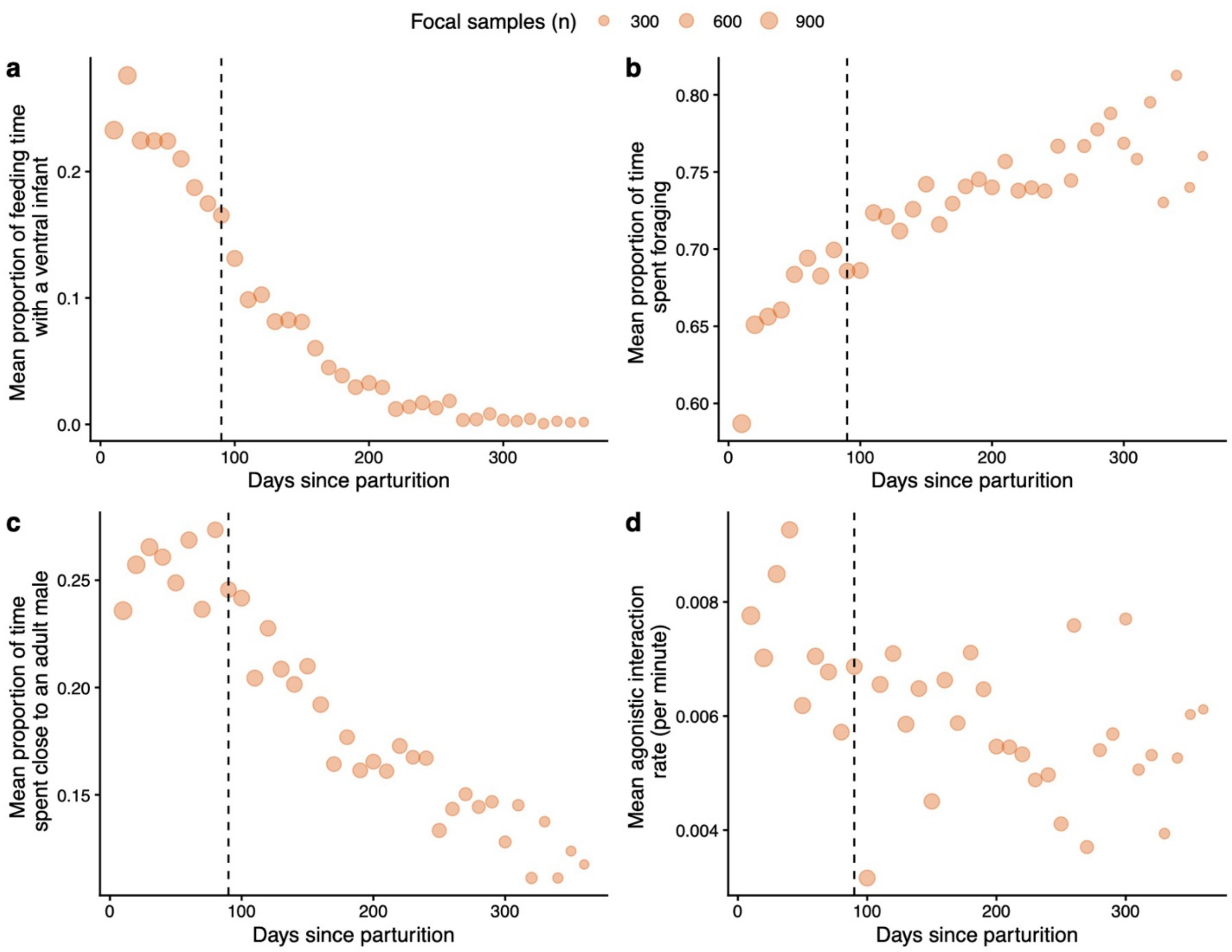

*Figure S2. The first 90 days postpartum are characterized by a dual challenge to mothers that affects both time budgets and behavior (10-day interval values).* As with daily values shown in Figure S1, data aggregated over 10-day intervals show that in the first 90 days postpartum mothers spent (a) a greater proportion of feeding time spent in ventral contact with an infant, (b) a lower proportion of time foraging, and (c) a greater proportion of time within 5 m of adult males, than females in later periods postpartum. (d) Female-female agonistic interaction rates were also higher when aggregated over 10-day intervals than during later postpartum periods. Each orange circle represents the mean value across all focal samples conducted in a given 10-day interval, scaled proportionally in size to the number of focal samples in our data set which were conducted in that interval. The dotted vertical line indicates 90 days postpartum, our defined threshold demarcating early lactation from the rest of postpartum amenorrhea.

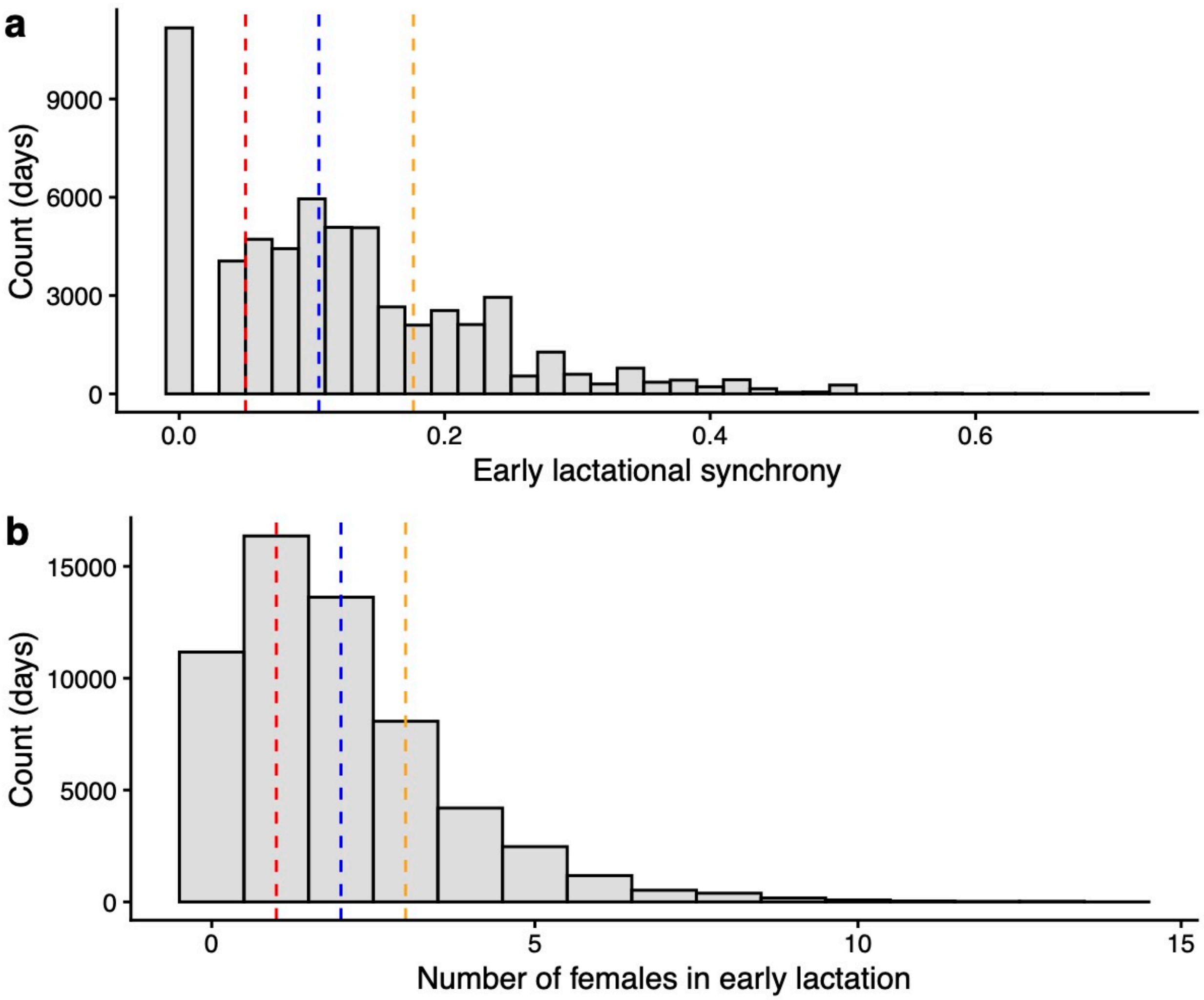

*Figure S3. Distribution of early lactational synchrony across the study period*. Histograms of daily values of (a) early lactational synchrony (the proportion of adult females in the group with an infant <90 days old), and (b) the number of females in the group who had an infant <90 days old from 1976-2021, calculated across 58,322 group-days for 19 distinct social groups (two original study groups and their descendent groups). The red, blue, orange dashed lines indicate the 25th, 50th, and 75th percentile values, respectively, in both panels.

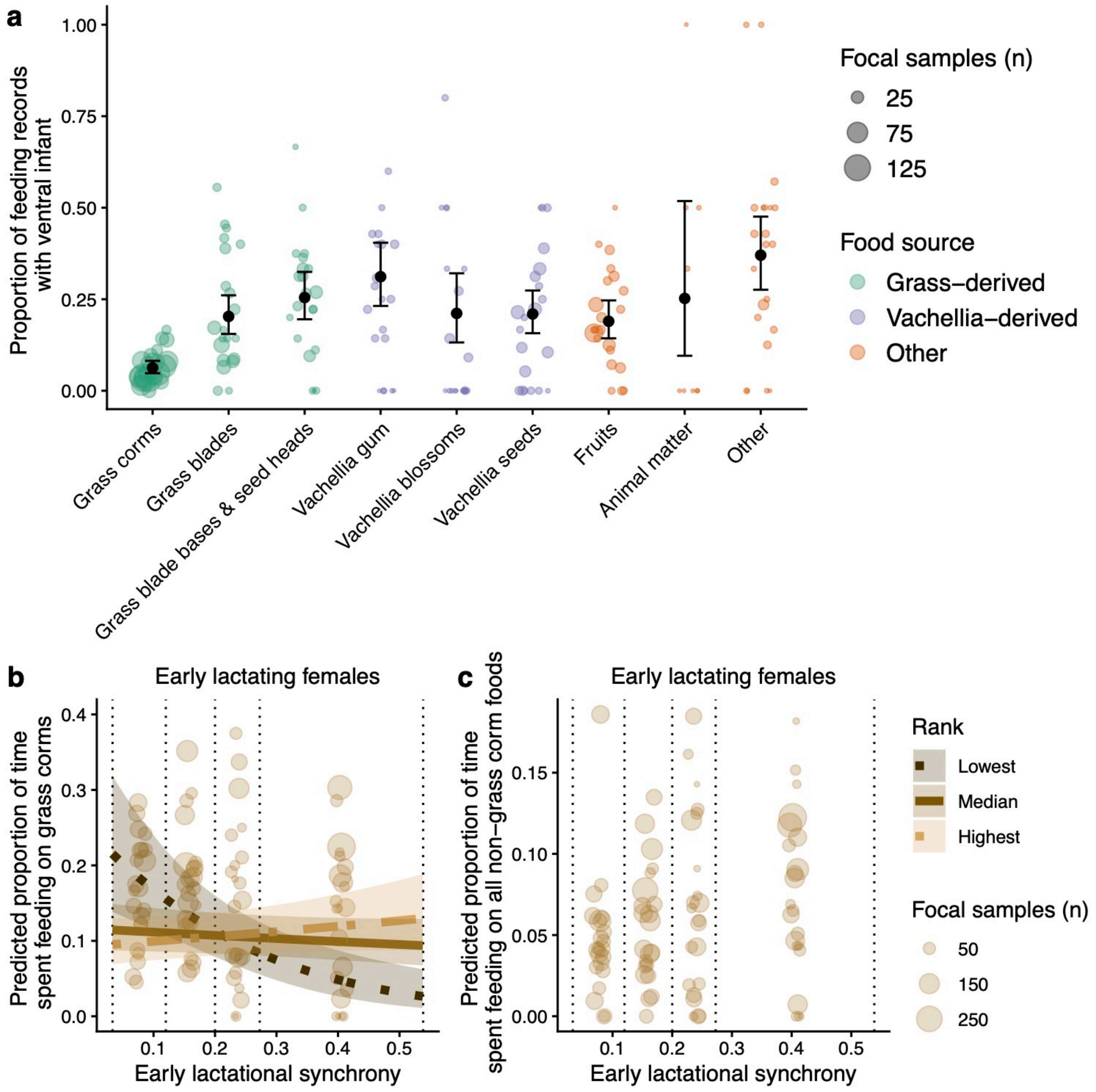

*Figure S4. Ventral infant carrying interferes with grass corm feeding*. (see full caption on next page).

*Figure S4. Ventral infant carrying interferes with grass corm feeding*. (a) Early lactating females were less likely to be in ventral contact with their infant when they were feeding on grass corms than on any other food category. The black circles represent GLMM-predicted values for the response variable across each food category (holding all other fixed effects constant at their observed mean value) and the error bars represent the 95% confidence intervals of these predicted values. The colored circles represent the observed mean probability in a complete hydrological year. (b) Low-ranking females, but not other females, decreased their proportion of total time spent feeding on grass corms as early lactational synchrony increased. The colored lines depict the GLMM-predicted relationship between early lactational synchrony and the response variable for alpha females (lightest-colored dot-dashed line; ordinal rank = 1), median-ranked females (solid line, ordinal rank = 7), and the lowest-ranked females in our dataset (darkest-colored dotted line, ordinal rank = 29). The shaded regions around these lines represent the 95% confidence intervals of these predicted relationships. The colored circles represent the mean observed proportions calculated from all data collected on days that were within one quantile of early lactational synchrony in a complete hydrological year. The vertical dotted lines indicate the quantiles of early lactational synchrony. (c) There was no significant relationship between rank, early lactational synchrony, and the proportion of total time spent eating foods other than grass corms. As in (c), the colored circles represent the mean observed proportions calculated from all data collected on days that were within one quantile of early lactational synchrony in a complete hydrological year and vertical dotted lines indicate the quantiles of early lactational synchrony.

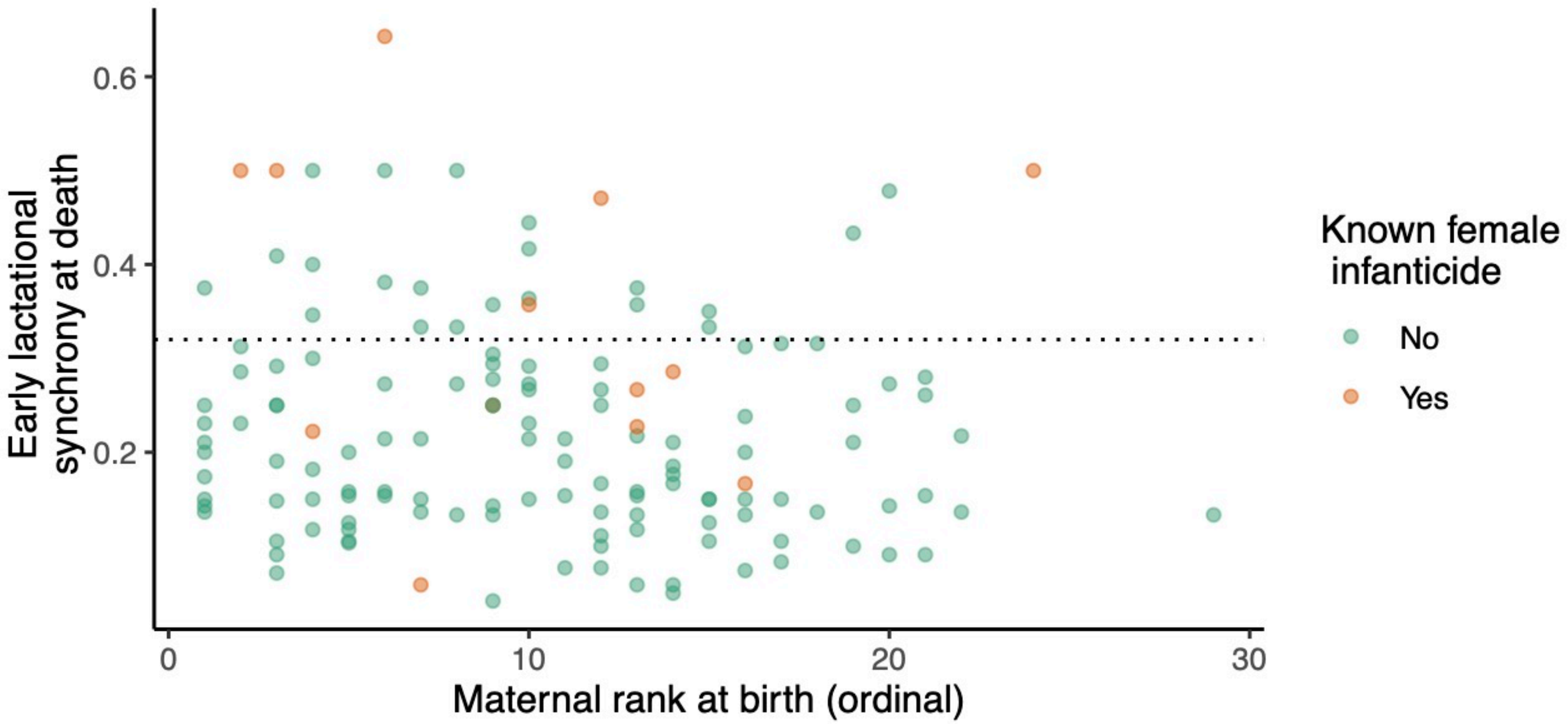

*Figure S5. Early lactational synchrony is a recurring feature of our population, and infant deaths to mothers of all ranks are well-distributed across values of early lactational synchrony.* Each circle represents one infant death in our sample (n = 140). Orange circles represent known or strongly suspected case of within-group infanticides by females, while green circles represent all other infant deaths. The majority of infant death causes were unknown. The dotted horizontal line is positioned at 0.320, the 95$^{th}$ percentile value of early lactational synchrony across all observed group-days.

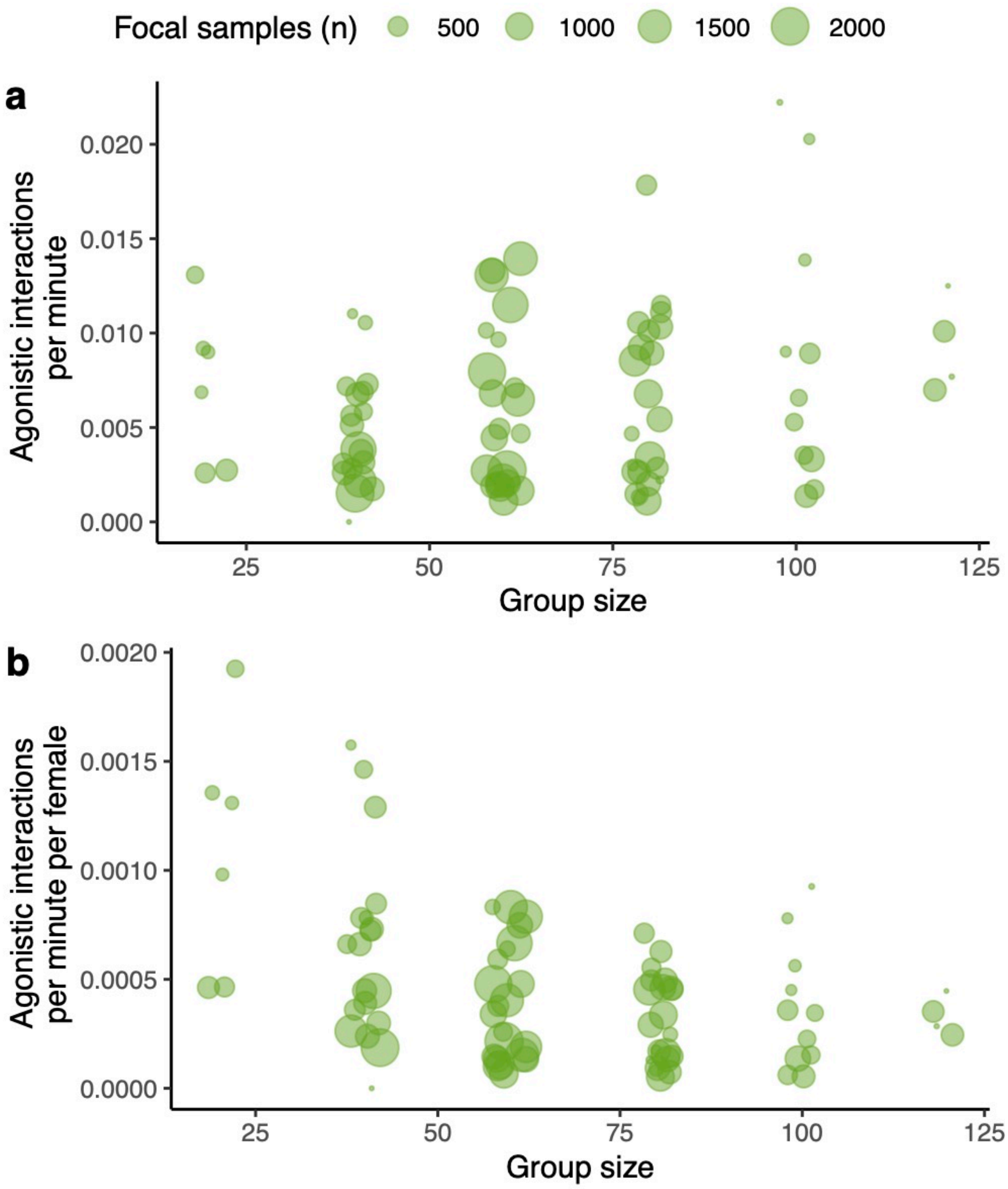

*Figure S6. Female-female agonistic interaction rate as a function of group size.* (a) depicts the observed relationship between total group size and the rate (per minute) at which focal adult females interacted agonistically with another adult female during focal sampling. (b) depicts this same relationship, but agonistic interactions rate divided by the number of nonfocal adult females who were in the group at the time of the focal sample. Each circle represents the mean value observed in one hydrological year and in one group size bin. Group sizes were binned into the following intervals: (0,20], [21,40], [41,60], [61,80], [81,100], [101,120].

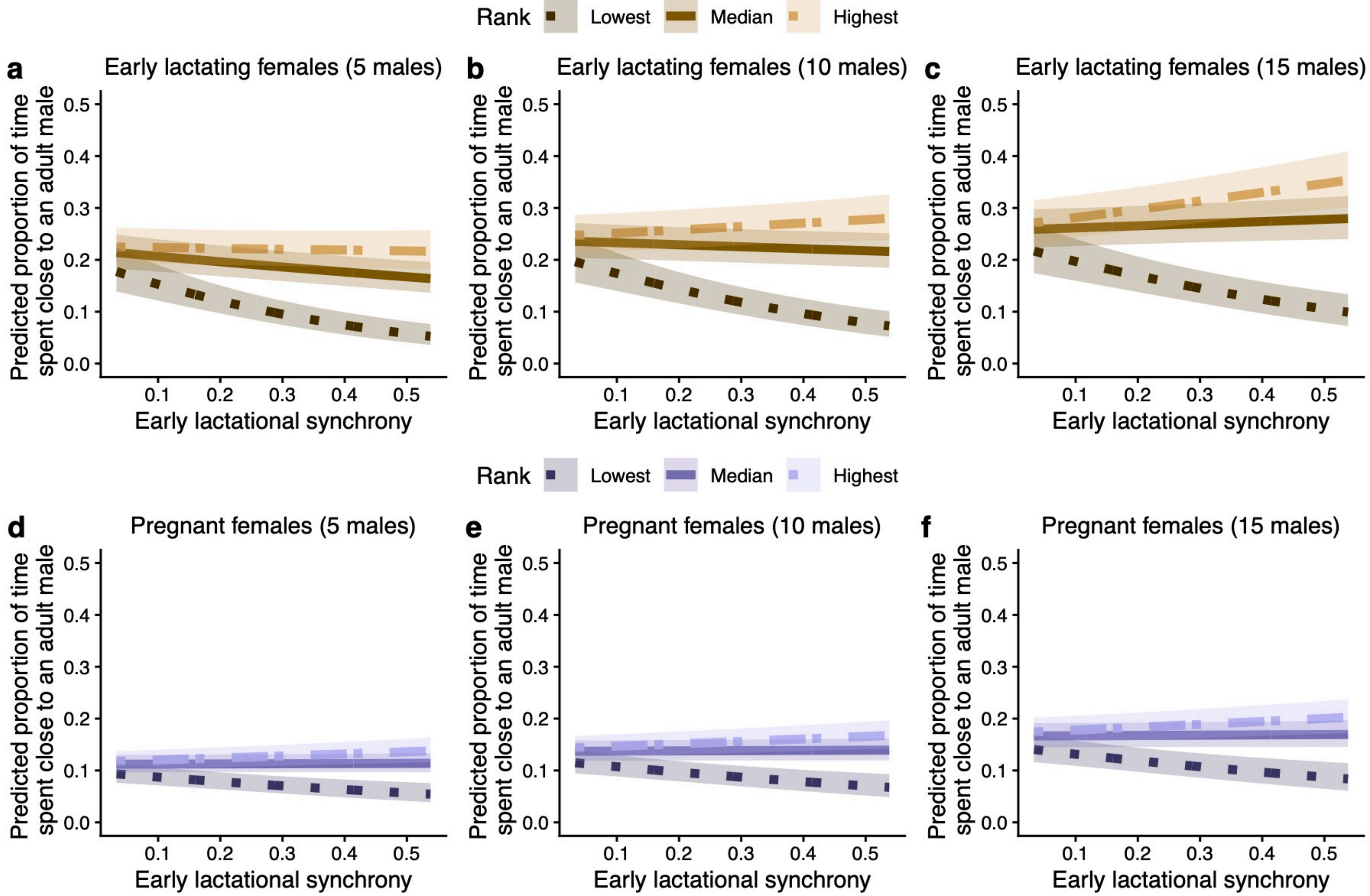

*Figure S7. The number of adult males, dominance rank, early lactational synchrony, and time spent with males*. The number of adult males in a group mediates the relationship between early lactational synchrony, dominance rank, and the proportion of time that (a-c) early lactating females spend with males, but not the proportion of time that (d-e) pregnant females spend with males. All panels depict the GLMM-predicted relationship between early lactational synchrony and the proportion of time spent with males for females of the lowest, median, and highest dominance rank when (a,d) five males, (b,e) 10 males, or (c,f) 15 males were present.

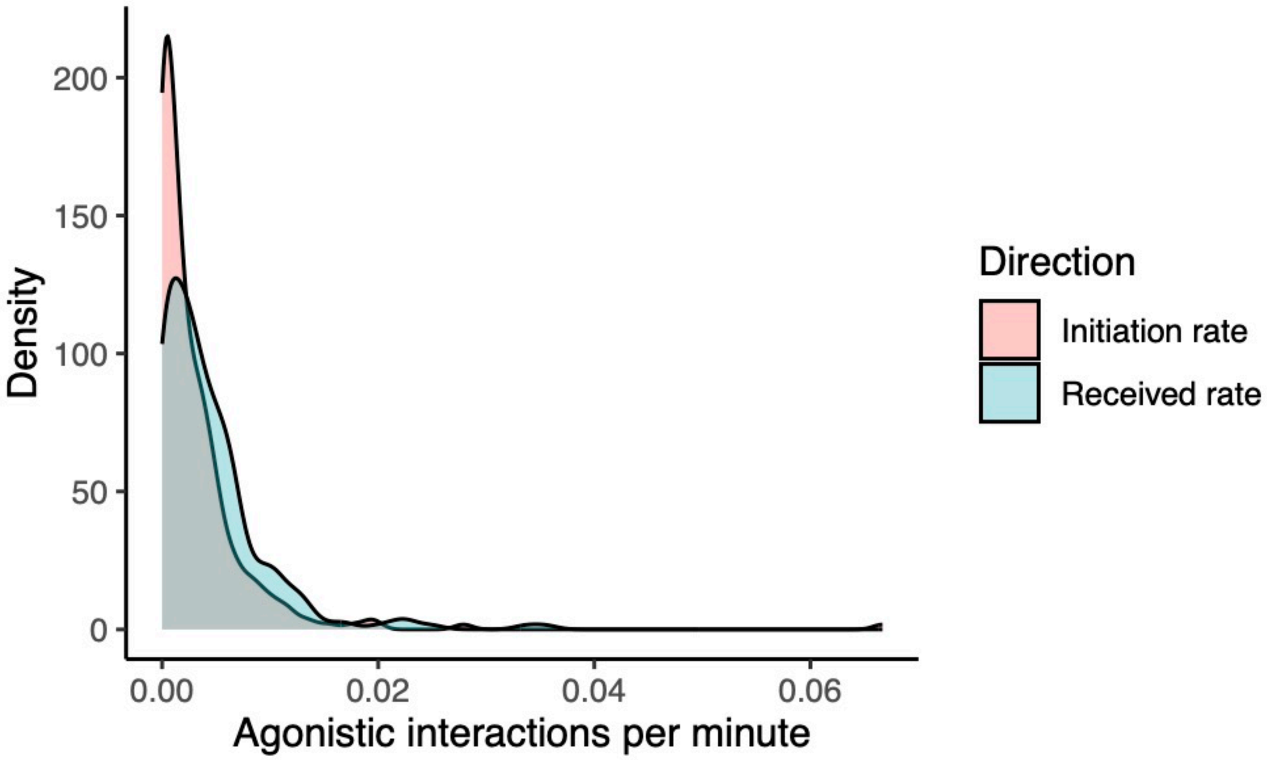

*Figure S8. Individual variation in rates of agonistic interactions initiated versus received.* Density plots illustrate the distribution of mean agonistic interaction rates calculated at the individual-level for adult females (n = 270 females), separated by interactions initiated by (red) versus received by (blue) focal females. Relatively more females exhibit low levels of interaction initiation than exhibit low levels of interaction reception, which contributes to lower mean rates of interaction initiation than of interaction reception when calculated across females.

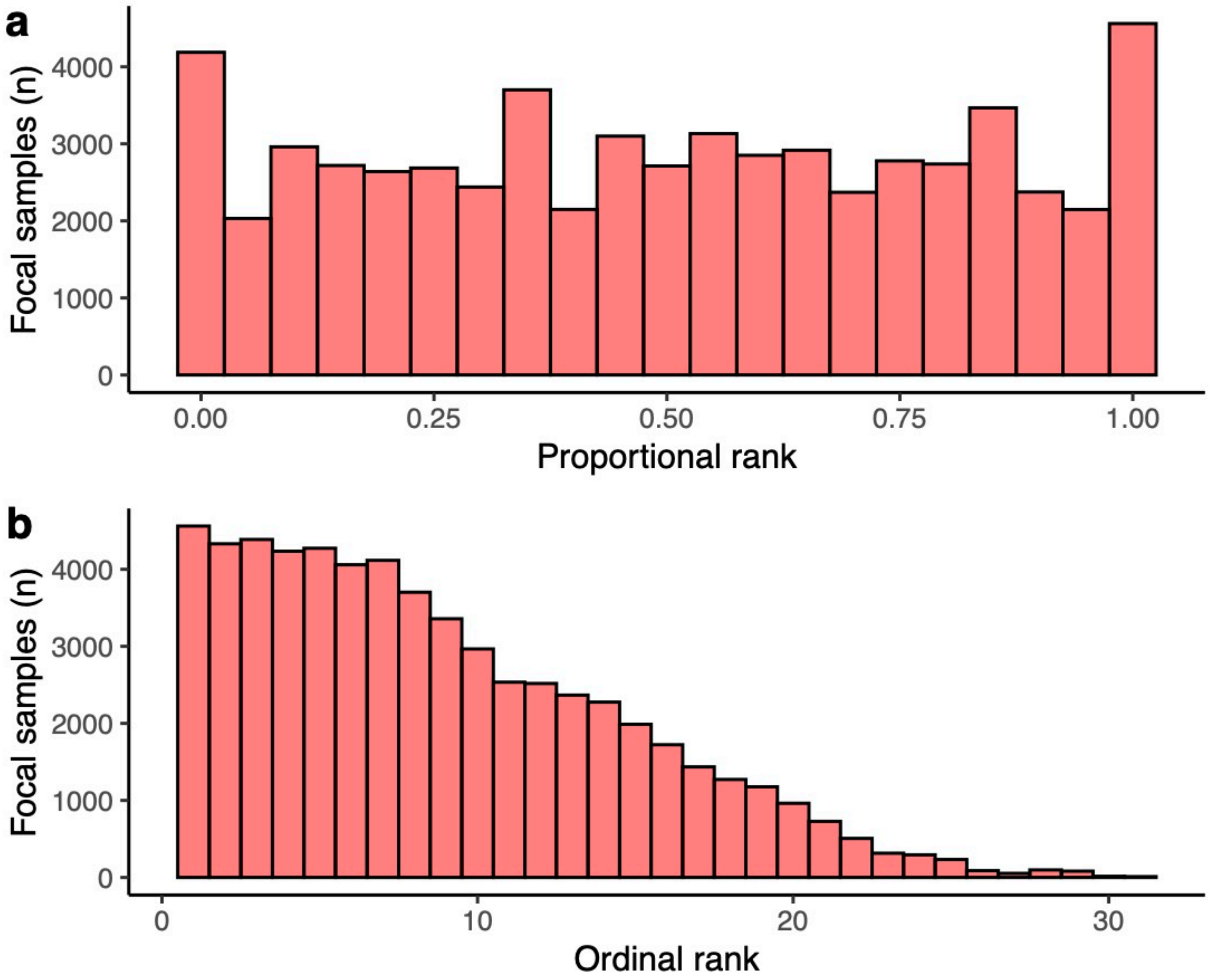

*Figure S9. Distributions of focal female dominance ranks during focal sampling.* (a) depicts the distribution of focal female proportional ranks at the time focal samples were conducted and (b) depicts the distribution of focal female ordinal ranks at the focal samples were conducted.
